# Arginine Metabolism Supports *De Novo* Pyrimidine Biosynthesis to Block DNA Damage and Maintain Epstein-Barr Virus Latency

**DOI:** 10.1101/2025.10.21.683727

**Authors:** Shaowen White, Yifei Liao, Eric M. Burton, John M. Asara, Benjamin E. Gewurz

**Affiliations:** Division of Infectious Diseases, Department of Medicine, Brigham and Women’s Hospital, 181, Longwood Avenue, Boston, MA 02115, USA; Broad Institute, Cambridge, MA 02142, USA; Division of Signal Transduction, Beth Israel Deaconess Medical Center and Department of Medicine, Boston, MA 02215, USA

**Keywords:** viral latency, lytic cycle, reactivation, metabolism, epigenetic, DNA damage, de novo pyrimidine synthesis, nucleotide biosynthesis, nucleotide metabolism, double-stranded DNA virus

## Abstract

Incompletely understood mechanisms serve to maintain Epstein-Barr virus (EBV) latency in most B-cell states, in which viral oncogene(s) are expressed but lytic antigens are repressed. Shortly after EBV’s discovery and even before it was named, early pioneers Werne and Gertrude Henle identified that restriction of extracellular arginine de-represses EBV lytic antigens within Burkitt lymphoma tumor cells. However, for nearly 60 years, it has remained unknown how arginine metabolism supports EBV latency. To gain insights, we performed an amino acid restriction screen in Burkitt cell lines. This confirmed that arginine restriction was sufficient to trigger EBV reactivation in Burkitt B-cells and gastric carcinoma models. Arginine restriction strongly impaired *de novo* pyrimidine biosynthesis, and CRISPR or chemical genetic blockade of pyrimidine biosynthesis enzymes induced EBV immediate early and early lytic gene expression. However, arginine restriction blocked EBV lytic DNA replication and consequently also late gene expression, suggesting an abortive lytic cycle. Arginine restriction triggered DNA damage, which was an important driver of arginine restriction-driven EBV reactivation. Arginine restriction and DNA hypomethylation synergistically increased EBV reactivation. Together, our results highlight arginine and pyrimidine metabolism as potential targets for EBV lytic antigen induction therapy in B and epithelial cell contexts.

**Importance:** Altered metabolism is a hallmark of cancer, frequently increasing transformed cell dependence on extracellular amino acid supply. Despite current interest in EBV lytic antigen induction therapy, in which viral lytic reactivation sensitizes tumors to the highly cytotoxic effects of the antiviral ganciclovir, there has been no systemic study of extracellular amino acid that controls EBV latency. We identified that arginine uptake was important for the maintenance of EBV latency in both Burkitt lymphoma and gastric carcinoma contexts. Metabolic pathway analyses highlighted that arginine uptake and metabolism was required to supply pyrimidines. Disruption of arginine metabolism or *de novo* pyrimidine synthesis caused DNA damage. As arginine restriction was also found to cause Burkitt DNA hypermethylation, we provide evidence that the combination of arginine restriction and DNA hypomethylation by decitabine or by CRISPR approaches together induced EBV reactivation more highly than either alone, suggesting a therapeutic approach.

## Introduction

The gamma-herpesvirus Epstein-Barr virus (EBV) chronically infects over 95% of adults worldwide (1). EBV was the first human tumor virus discovered through its close association with the endemic Burkitt lymphoma (BL), which remains the most common pediatric non-Hodgkin B cell lymphoma in areas with holoendemic malaria (2, 3) EBV contributes to ∼1.5% of all human cancers, including multiple subtypes of both Hodgkin and non-Hodgkin B-cell lymphomas, natural killer and T cell lymphomas, gastric and nasopharyngeal carcinoma (2, 4–7). In most infected tumor cells, EBV maintains a state of viral latency, in which nearly 80 viral lytic cycle genes are repressed.

EBV infects both lymphocytes and epithelial cells to persistently infect hosts and spread between individuals. EBV uses a series of latency genes to traverse the B cell compartment and colonize the memory B cell reservoir (8). EBV latency programs, which express between 1-9 viral oncoproteins and EBV non-coding RNAs, expand the infected B-cell pool and enable infected cells to differentiate into long-lived memory B-cells (8–10). Within latently infected cells, the double-stranded DNA viral genome is maintained as circular, extrachromosomal episomes.

Memory B-cell differentiation into plasma cells is thought to be the major physiological B-cell trigger for EBV reactivation. Within Burkitt tumor cells, which are a major model for studies of EBV reactivation as robust primary cell models of EBV reactivation have yet to developed, B-cell immunoglobulin receptor cross-linking triggers reactivation, as does treatment with tumor promoting agent (TPA, also called phorbol myristate acetate) and the histone deacetylase inhibitor sodium butyrate (11, 12). Many EBV-infected tumors cells, including Burkitt and gastric carcinoma cells, exhibit high levels of DNA CpG hypermethylation, which support viral latency. Upon lytic reactivation, the EBV-encoded immediate early genes BZLF1 and BRLF1 are first to be expressed, which are transcription activators that induce expression of ∼35 early lytic viral genes (13, 14). These include factors required for lytic cycle EBV genome amplification, including the processivity factor BMRF1 and the protein kinase BGLF4 (15–20). Newly synthesized, linear, unchromatinized viral genomes serve as the templates for expression of ∼37 EBV late genes (21–23), which include virion structural proteins including the capsid protein p18 and virion glycoproteins, including glycoprotein 350 (gp350) (24, 25). BGLF4 sensitizes lytic cells and their neighbors to highly cytotoxic activity of the antiviral nucleoside ganciclovir (26, 27). EBV lytic antigens also promote potent innate and adaptive immune responses (15, 28, 29). Consequently, there is significant interest in development of strategies to induce EBV lytic reactivation to sensitize EBV+ tumors to ganciclovir and immunotherapy approaches (30–32).

Altered metabolism is a hallmark of cancer (33). In fact, Burkitt B-cells are amongst the fastest growing human tumor (34). Cancer cells frequently rewire major central carbon metabolism pathway to support aerobic glycolysis (the so-called Warburg effect) (35), increased dependency on *de novo* nucleotide biosynthesis (36), extracellular arginine (37), asparagine (38), cysteine (39) and methionine (40). Shortly after EBV was discovered but before it was named by Werner and Gertrude Henle, they serendipitously discovered that restriction of extracellular arginine de-repressed several EBV lytic antigens against which antisera was already available in the newly available human tumor derived Burkitt cell lines (41). However, the mechanism was not pursued at that early period of Burkitt tumor research and key questions have remained unaddressed, such as to what extent and how does arginine restriction (AR) induce EBV lytic antigen expression and in which latently infected host cellular contexts does this phenomenon occur.

Arginine is a conditionally essential amino acid whose *de novo* synthesis in small intestine and kidney meets the need in healthy adults, but is essential (must be acquired by diet) in infants, growing children, and adults under catabolic stress or with dysfunction of the small intestine or kidney (42, 43). Its metabolism is a vulnerability of many types of cancer cells, especially in which the gene encoding argininosuccinate synthase (ASS1), the rate-limiting enzyme in arginine biosynthesis pathway (44, 45), has been epigenetically silenced. In addition to its obligatory role in protein biosynthesis, arginine metabolism yields a range of important metabolites (46), which serve as major regulators of mitochondria physiology, including respiration and mitochondrial biosynthesis (47–49), and nucleotide metabolism. Arginine supports, eukaryotic pyrimidine *de novo* biosynthesis supports growth and proliferation of EBV-transformed B-cells (50). Notably, halogenated pyrimidine analogs such as 5-fluorouracil and idoxuridine and the antifolate methotrexate each trigger EBV lytic gene expression (51–54). Methotrexate disrupts tetrahydrofolate synthesis, which is required for *de novo* purine biosynthesis (55). These observations suggest that nucleotide metabolism supports EBV latency through incompletely defined mechanisms.

The *de novo* pyrimidine biosynthesis pathway relies on a series of enzymes including the rate limiting CAD (carbamoyl-phosphate synthetase II, aspartate trans-carbamoylase and dihydroorotase), UMPS (uridine 5′-monophosphate synthase) and the mitochondrial membrane protein DHODH (dihydroorotate dehydrogenase). CAD mediates the biosynthesis of dihydroorotate from carbamoyl-phosphate and aspartate. DHODH and UMPS then mediate downstream biosynthesis of uridine 5’-monophosphate (UMP). These proteins form a complex called the pyrimidinosome which spans across the inner and outer mitochondrial membranes (56). Pyrimidinosome activity is regulated by AMP-activated protein kinase (AMPK), whose activity is in turn regulated by the AMP/ATP and ADP/ATP ratios (56, 57) or by mitochondrial electron transport chain activity (58). In bacteria, *de novo* pyrimidine synthesis activity is regulated by arginine levels, as both arginine and pyrimidine biosynthesis consume carbamoyl phosphate (59), suggesting that arginine and pyrimidine biosynthesis are evolutionarily related pathways.

To systematically identify amino acids that contribute to EBV latency, we first conducted an amino acid restriction screen, which confirmed arginine’s essential roles in EBV latency maintenance. Metabolomic analyses revealed that extracellular arginine is required for Burkitt pyrimidine *de novo* biosynthesis. Restriction of extracellular arginine levels disrupted *de novo* pyrimidine biosynthesis at the level of aspartate transcarbamoylation, and perturbation of *de novo* pyrimidine biosynthesis therefore also triggered EBV lytic reactivation. Arginine restriction-mediated EBV reactivation required host cell DNA damage response pathways and synergized with DNA hypomethylation agents to further drive EBV lytic antigen de-repression.

## Results

### An amino acid restriction screen for Burkitt lymphoma EBV lytic reactivation

To systematically test whether supply of particular amino acid roles was necessary for EBV latency, we performed an amino acid restriction screen in P3HR-1 Burkitt lymphoma cells, using a panel of RPMI media lacking one of each of the 20 amino acids, together with fetal calf serum (FCS) dialyzed to remove free amino acids (**Figure 1B**). Restriction of any of the nine essential amino acids led to cell growth arrest (**Figure S1A**). Restriction of non-essential amino acids had comparatively little effect on Burkitt proliferation, as expected, with the exception of arginine, cystine (the oxidized form of cysteine present in media), glutamine and tyrosine (**Figure S1B**). Cystine restriction resulted in widespread cell death by day 3, likely due to its obligatory roles in redox defense against lipid peroxidation (60). Thus, cystine restriction was omitted from further analyses.

**Figure 1.**
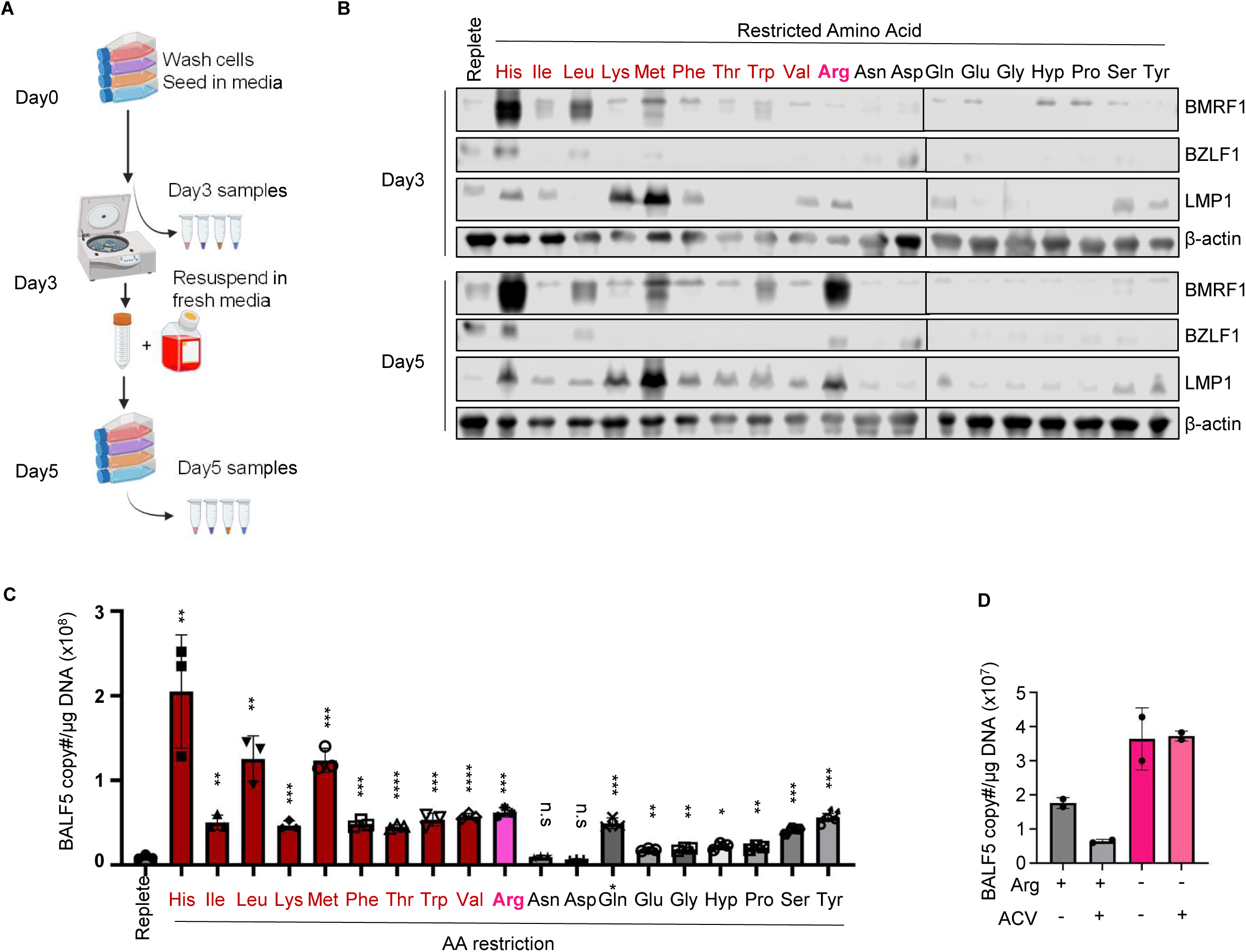
Screen of amino acid restriction effects on Burkitt B cell EBV reactivation. (A) Schematic of amino acid restriction of P3HR-1 Burkitt cells. (B) Immunoblot analysis of whole-cell lysates (WCL) from P3HR-1 grown in replete media vs in media restricted for the indicated amino acid for 3 or 5 days. Blots are representative of n=2 replicates. (C) Analysis of amino acid restriction effects on P3HR-1 EBV genome copy number. Shown are the mean ± SEM EBV genome copy number as determined by qPCR analysis from n=3 replicates. (D) Mean EBV intracellular genome copy number from qPCR analysis of n=2 biological replicates, each with two technical replicates, of P3HR-1 grown in replete or arginine-free media for 5 days, with our without 100 mg/ml acyclovir (ACV) to block EBV lytic replication. Student’s T-test was performed, with ****p < 0.0001, *p < 0.05 compared to replete control media.

To define amino acid restriction effects on EBV latency, immunoblot analysis was performed on whole cell lysates of P3HR-1 following three or five days of incubation with media restricted for one of each amino acid. Restriction of methionine or lysine de-repressed the EBV oncoprotein LMP1, which is expressed by the EBV latency IIa or III programs and also by the lytic cycle, and to a lesser degree the EBV immediate early lytic BZLF1 and early lytic BMRF1 proteins. By contrast, restriction of arginine, histidine or leucine more highly de-repressed BZLF1 and BMRF1, particularly by day 5 (**Figure 1B**). While amino acid restriction can inhibit the metabolism master regulator mTOR, inhibition of mTOR itself by rapamycin did not de-repress EBV lytic antigen expression (**Supplemental Figure S1C**), suggesting that other metabolic pathway(s) underlie maintenance of EBV latency.

To next define amino acid restriction effects on EBV lytic genome replication, which is carried out in the early lytic phase by a viral DNA polymerase, we performed qPCR analysis of EBV genome copy number. Surprisingly, restriction of any of the essential amino acids, or of several non-essential amino acids, increased EBV genome copy number to varying degrees, which could be secondary to effects on replication of either latent or lytic EBC genomes. Of these, histidine, leucine, methionine and arginine restriction most highly increased EBV genome copy number in comparison to levels observed in Burkitt cells grown in replete media (**Figure 1C**). Since EBV genome copy number can also differ within latency (61), to differentiate between effects on latent versus lytic genome replication, we repeated the assay in the presence of acyclovir to block lytic EBV genome amplification. Interestingly, arginine restriction increased EBV copy # even in acyclovir treated cells (**Figure 1D** and **S1D**). Therefore, arginine restriction may amplify latent EBV genome copy number either by provoking extra rounds of latent episome replication versus increased levels of replication within the S-phase (62, 63).

### Arginine restriction in culture media leads to abortive lytic reactivation

Although histidine restriction highly induced EBV reactivation in P3HR-1, it did not induce EBV lytic antigen in two other Burkitt cell lines (**Figure S1E**), including the EB3 Burkitt lymphoma cells used by the Henles in their seminal arginine restriction study. Since we observed strong arginine restriction effects across Burkitt models, we focused on how it de-represses immediate early and early lytic EBV antigens. First, to define the extent of arginine restriction required for EBV reactivation, Burkitt cells were incubated in media with a range of arginine concentrations. RPMI contains 1.5 mM arginine, whereas plasma arginine typically ranges from 41μM-114 μM, and can be lowered by 50% by dietary arginine restriction to the 20-70μM range (64, 65). Reduction of arginine RPMI levels by 90% to 115μM, similar to the physiological arginine level, did not alter Burkitt cell growth over 7 days (**Figure S2A**). By contrast, further restriction of extracellular arginine by 99% to 11.5 μM induced growth arrest, but only low levels of cell death. Similar observations even extended to Burkitt cells cultured in arginine-free media for 5 days (**Figure S2B**). In cells cultured for 5 days at 11.5μM (1% of typical RPMI arginine concentration), BZLF1 and BMRF1 were de-repressed, which was also observed in the human tumor derived EBV+ gastric cancer cell line SNU-719 (**Figure 2A-B**). BMRF1 expression was also readily detectable by confocal immunofluorescence microscopy analysis in cells cultured in arginine-free media for 5 days (**Figure 2C-D**).

**Figure 2.**
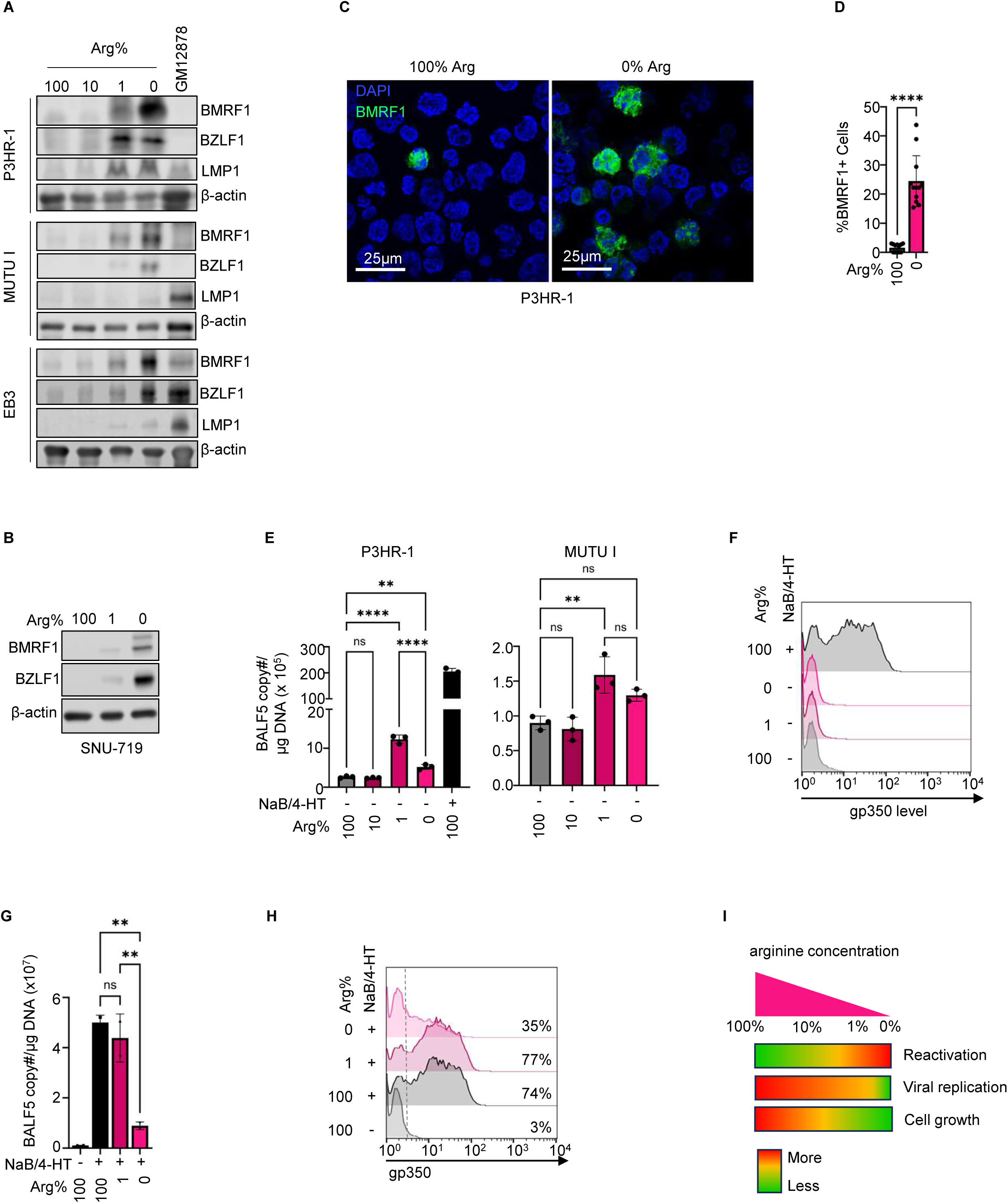
Characterization of arginine restriction effects on EBV lytic reactivation. (A) Analysis of arginine restriction effects on BZLF1 and LMP1 de-repression. Immunoblot analysis of WCL from P3HR-1, MUTU I or EB3 Burkitt cells cultured in media with the indicated arginine level for 5 days. (B) Arginine restriction effects on EBV reactivation in SNU-719 gastric carcinoma cells. Immunoblot of WCL from SNU-719 cultured in RPMI with the indicated Arg level for 5 days. (C) Confocal immunofluorescence analysis of BMRF1 expression in P3HR-1 cells cultured in replete or arginine-free media for 5 days. (D) Mean ± SD percentages of BMRF1+ cells from n=13 fields as in (C). (E) qPCR analysis of intracellular EBV genome copy number in P3HR-1 or MUTU I Burkitt cells cultured in media with the indicated arginine concentration for 5 days. As a positive control, P3HR-1 with a conditional 4-hydroxytamoxifen (4-HT) responsive lytic reactivation system (ZHT/RHT) grown in replete media and reactivated with 4-HT (400 nM) and sodium butyrate (NaB, 0.5 mM) for 24 hours are shown. Data are shown as BALF5 copy number per mg of DNA. (F) FACS analysis of plasma membrane gp350 levels in P3HR-1 cells treated as in (E). (G) qPCR analysis of EBV genome copy number in P3HR-1 ZHT/RHT cells cultured in media with the indicated arginine concentrations and reactivated by 4HT/Nab treatment for 24 hours. Shown are mean values from n=2 replicates, each with two technical replicates. (H) FACS analysis of plasma membrane gp350 values in P3HR1 ZHT/RHT cells treated as in (G). (I) Summary of extracellular arginine effects on EBV reactivation. Red represents stronger reactivation, green represents stronger maintenance of latency. Two-way ANOVA, one-way ANOVA, or Student’s t test was performed, with ****p < 0.0001, ***p < 0.001, **p < 0.01, *p < 0.05. Blots in (A) and (B) are representative of n=2 replicates.

When cultured in media with 1% or 0% the levels of arginine found in RPMI for 5 days together with dialyzed fetal calf serum to remove free amino acids, LMP1 expression was also de-repressed in P3HR-1 cells. LMP1 de-repression was observed to a lesser degree in EB3, but not in MUTU I Burkitt cells (**Figure 2A**). Since LMP1 can be expressed as a latency oncogene within the latency IIa or III programs versus as a lytic gene whose expression is highest at late periods of reactivation with roles that both support and inhibit the lytic cycle (66–68), it is possible that such complex regulation accounts for differences across Burkitt lymphoma models.

We next assed whether arginine restriction triggered viral lytic cycle DNA replication and the expression of EBV late antigen gp350, which is expressed from newly synthesized lytic EBV genomes. Arginine restriction mildly increased EBV genome copy number, though gp350 expression was not detected (**Figure 2E-F**). Interestingly, although arginine-free media more highly de-repressed BZLF1 and BMRF1 than culture in 1% of typical RPMI arginine levels, EBV genome copy number was significantly higher when cells were induced by 4-HT/NaB for lytic reactivation in media with 1% RPMI arginine levels than in arginine-free media (**Figure 2G**).

We therefore hypothesized that arginine might be essential for EBV lytic DNA replication, as arginine is a key building block for nucleotide synthesis and is important for herpes simplex virus replication (69). To test this hypothesis, P3HR-1 cells that express a conditional lytic reactivation system, in which tamoxifen (4-HT) triggers nuclear translocation of BZLF1 and BRLF1 immediate early proteins (70, 71), were seeded in media with a range of arginine concentrations and induced for lytic reactivation by 4HT and the histone deacetylase inhibitor sodium butyrate (NaB). Viral DNA replication and late protein gp350 expression were only modestly decreased when cells were grown in media containing 1% of typical RPMI arginine concentration. By contrast, culture in arginine-free media strongly blocked EBV lytic DNA replication and late lytic protein gp350 expression (**Figure 2G-H)**. These data indicate that Burkitt cell proliferation and viral lytic DNA replication differ in their dependence on extracellular arginine concentration (**Figure 2I**), and that severe arginine restriction resulted in abortive EBV reactivation, where immediate early and early genes are de-repressed, but viral lytic genomes are not synthesized and consequently late genes are not expressed.

### Metabolomic analysis of arginine restricted Burkitt cells

To further investigate metabolic signals and pathways that underlie arginine restriction-driven EBV reactivation, we performed liquid chromatography-mass spectrometry (LC-MS) analyses of polar metabolites, using extracts from EB3 Burkitt cells incubated in replete RPMI/FCS versus arginine-free RPMI/FCS for 5 days. As expected, arginine itself, and several closely related arginine metabolites, including ornithine, citrulline, and argininosuccinate were highly depleted in cells grown in arginine-free media. Interestingly, multiple nucleosides and nucleotide precursors were also significantly depleted by arginine restriction, while the pyrimidine precursor carbamoyl phosphate was instead more abundant in cells grown in arginine-free media (**Figure 3A**).

**Figure 3.**
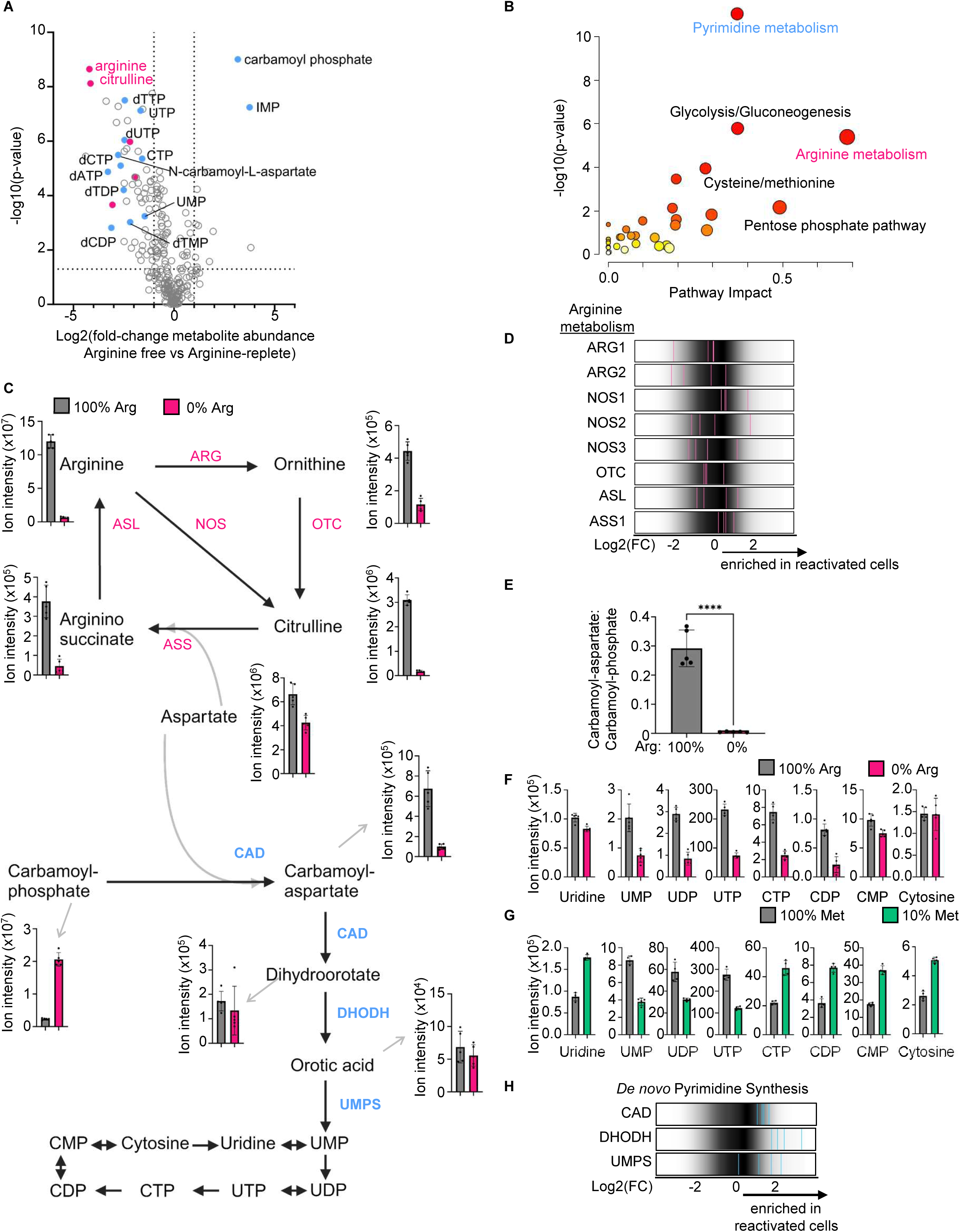
Arginine restriction impairs Burkitt d*e novo* pyrimidine biosynthesis. (A) Analysis of arginine restriction effects on the Burkitt B cell metabolome. Volcano plot of LC/MS metabolomic analysis of EB3 Burkitt cells cultured arginine free vs arginine-replete media for 5 days, from n=5 replicates. Higher foldchange values indicate higher levels in cells grown under arginine free conditions. Arginine cycle metabolites are highlighted in pink; pyrimidine and purine related metabolites are highlighted in blue. (B) KEGG metabolic pathway analysis of LC/MS data as in (A). Metabolites were selected by FDR < 0.05 and pathway impact values were computed by MetaboAnalyst 3.0 topological analysis. (C) Schematic of arginine cycle and *de novo* pyrimidine synthesis pathways. Shown at right are abundances of indicated metabolites in EB3 cells grown under arginine replete (100% Arg) versus Arginine free (0% Arg). (D) Rug plots of sgRNA levels targeting arginine metabolism genes from a human genome-wide CRISPR screen for host factors that maintain EBV latency in P3HR-1 cells (73). Shown are the Log2 transformed Foldchange abundances in gp350+ sorted P3HR-1 versus in the input library, where the magenta lines represent values for each of the four independent sgRNA targeting the specified gene. The overall distribution of human genomewide Brunello library sgRNAs are shown in gray in the background for reference, where most sgRNAs did not perturb EBV latency. Higher values indicate sgRNA enrichment in reactivated gp350+ cells versus in the input library as a whole. (E) CAD activity, as read out by the ratio of the CAD product carbamoyl-aspartate to the CAD precursor carbamoyl-phosphate in EB3 cells, as in (A). P-values were calculated by Student’s t-test. ****P < 0.0001. (F) Pyrimidine abundance from EB3 cells as in (A). (G) Abundance of pyrimidine metabolites from P3HR-1 cells cultured in 10% methionine vs 100% methionine media for 72 (74). (H) Rug plots highlighting foldchange abundances of the indicated *de novo* pyrimidine synthesis gene targeting sgRNAs in gp350+ cells shown in teal (four independent sgRNAs per gene) in a human genome-wide CRISPR screen for P3HR-1 factors that maintain EBV latency, as in (D)(73).

Metabolomic pathway analysis further highlighted that arginine and pyrimidine metabolism were among the most significantly altered processes in cells grown under arginine-free conditions (**Figure 3B**). To a lesser degree, glycolysis and gluconeogenesis pathways were also altered, consistent with prior reports that arginine starvation causes respiratory stress and inhibits aerobic glycolysis in transformed cells (47). While ATP levels were lower in cells grown under arginine-free conditions, perhaps reflecting growth-arrest, the ADP/ATP ratios were similar in cells grown under arginine-replete vs arginine-free conditions, suggesting that arginine restriction does not block energy production per se (**Figure S3A-B**).

The rate-limiting *de novo* arginine biosynthesis enzyme ASS1 combines citrulline and aspartate to produce argininosuccinate, which is then used by argininosuccinate lyase (ASL) to regenerate arginine (**Figure 3C**). Lymphomas frequently silence ASS1 expression (72), and such arginine auxotrophy can be leveraged by arginine restriction therapeutic approaches in pre-clinical development to trigger oxidative stress and mitochondria dysfunction (47, 49). We therefore surveyed ASS1 expression in EBV-infected Burkitt and gastric carcinoma cell contexts. Arginine restriction increased ASS1 expression in EB3 Burkitt cells and in SNU-719 gastric carcinoma cells, but not in two other Burkitt cell lines(**Figure S3C**), consistent with the variable ASS1 silencing across non-Hodgkin’s lymphomas (72).

We next examined if arginine restriction driven changes in arginine metabolite abundance might trigger EBV reactivation. Intracellular arginine can be catabolized by nitroxide (NO) synthases (NOS), arginine:glycine amidinotransferase, arginase (ARG) or by arginine decarboxylase to produce NO/citrulline, guanidinoacetate, ornithine and agmatine, respectively (**Figure 3C**). Ornithine can be further metabolized to generate citrulline via ornithine transcarbamylase (OTC). Since ornithine and citrulline were both significantly decreased by Burkitt cell culture in arginine-free media (**Figure 3C**), we investigated whether depletion of ARG, OTC, or NOS altered EBV reactivation. To gauge the involvement of these in EBV latency maintenance, we examined our lab’s recent human genome-wide CRISPR-Cas9 screen for host factors that maintain Burkitt EBV latency (73). CRISPR single-guide RNAs (sgRNAs) targeting arginine metabolism genes, including ARG, NOS or OTC were not enriched in cells triggered for EBV lytic reactivation by CRISPR editing, arguing against their obligatory roles in maintenance of EBV latency (**Figure 3D**).

We then examined changes in pyrimidine biosynthesis pathways in arginine restricted cells. *De novo* pyrimidine biosynthesis requires the trifunctional enzyme CAD (Carbamoyl-phosphate synthetase 2, aspartate transcarbamylase and dihydroorotase), which first uses its aspartate transcarbamoylase domain to synthesize carbamoyl-aspartate from carbamoyl-phosphate and aspartate (**Figure 3C**). The CAD hydroorotase domain then metabolizes carbamoyl-aspartate to dihydroorotate, which is converted to uracil monophosphate (UMP) by the enzymes dihydroorotate dehydrogenase (DHODH) and UMP synthase (UMPS) (56) (**Figure 3C**). Metabolomic analysis highlighted that CAD carbamoyl-phosphate synthetase activity, defined by the carbamoyl-phosphate to carbamoyl-aspartate precursor/product ratio, was strongly inhibited by arginine restriction, which correlated with decreased pyrimidine levels, including of UTP and CTP (**Figure 3E-F**). By contrast, Burkitt methionine restriction (74) instead increases CTP levels (**Figure 3G**). Intriguingly, CRISPR screen analysis suggests that perturbation of the *de novo* pyrimidine pathway by knockout of CAD, DHODH or UMPS triggers EBV reactivation (**Figure 3H**), as sgRNAs targeting each of these were highly enriched in the population in which CRISPR knockout induced EBV reactivation. Taken together, these data suggest that *de novo* pyrimidine synthesis and likely therefore also abundant pyrimidine supply supports maintenance of EBV latency.

### D*e novo* pyrimidine biosynthesis is necessary for Burkitt EBV latency

We next tested whether *de novo* pyrimidine biosynthesis was necessary for maintenance of EBV latency within Burkitt B cells. First, we used CRISPR to deplete CAD within P3HR-1 Burkitt cells (**Figure 4A**). We then performed LC/MS metabolomic analysis to define CAD KO effects on metabolite abundances. Suggestive of on-target CRISPR effects, the CAD precursor carbamoyl phosphate was the metabolite most highly increased by CAD depletion. Similar to arginine restriction, CAD depletion highly altered levels of multiple nucleotides and nucleotide-related metabolites, with UTP amongst the most decreased (**Figure 4B**). Similarly, shifting P3HR-1 from arginine replete to arginine-free media for five days strongly decreased the ratio of CAD product carbamoyl-aspartate to CAD precursor carbamoyl-phosphate and perturbed levels of multiple nucleotides (**Figure 4C-D**). Arginine restriction also highly depleted arginine the arginine metabolites citrulline and ornithine. Metabolomic pathway impact analysis highlighted that arginine restriction most highly perturbed arginine biosynthesis, as expected, but also highly perturbed purine and pyrimidine metabolism (**Figure 4D and S4A-B**). Arginine restriction similarly altered metabolites in P3HR-1 and EB3 Burkitt cells, in both of whom arginine, citrulline and ornithine levels were strongly diminished (**Figure S4C**). Interestingly, arginine restriction, but not CAD KO, strongly decreased intracellular arginine and citrulline levels (**Figure 4D**). Arginine restriction and CAD KO each depleted ATP, CTP and UTP levels, but increased TMP and TDP levels (**Figure 4D**).

**Figure 4.**
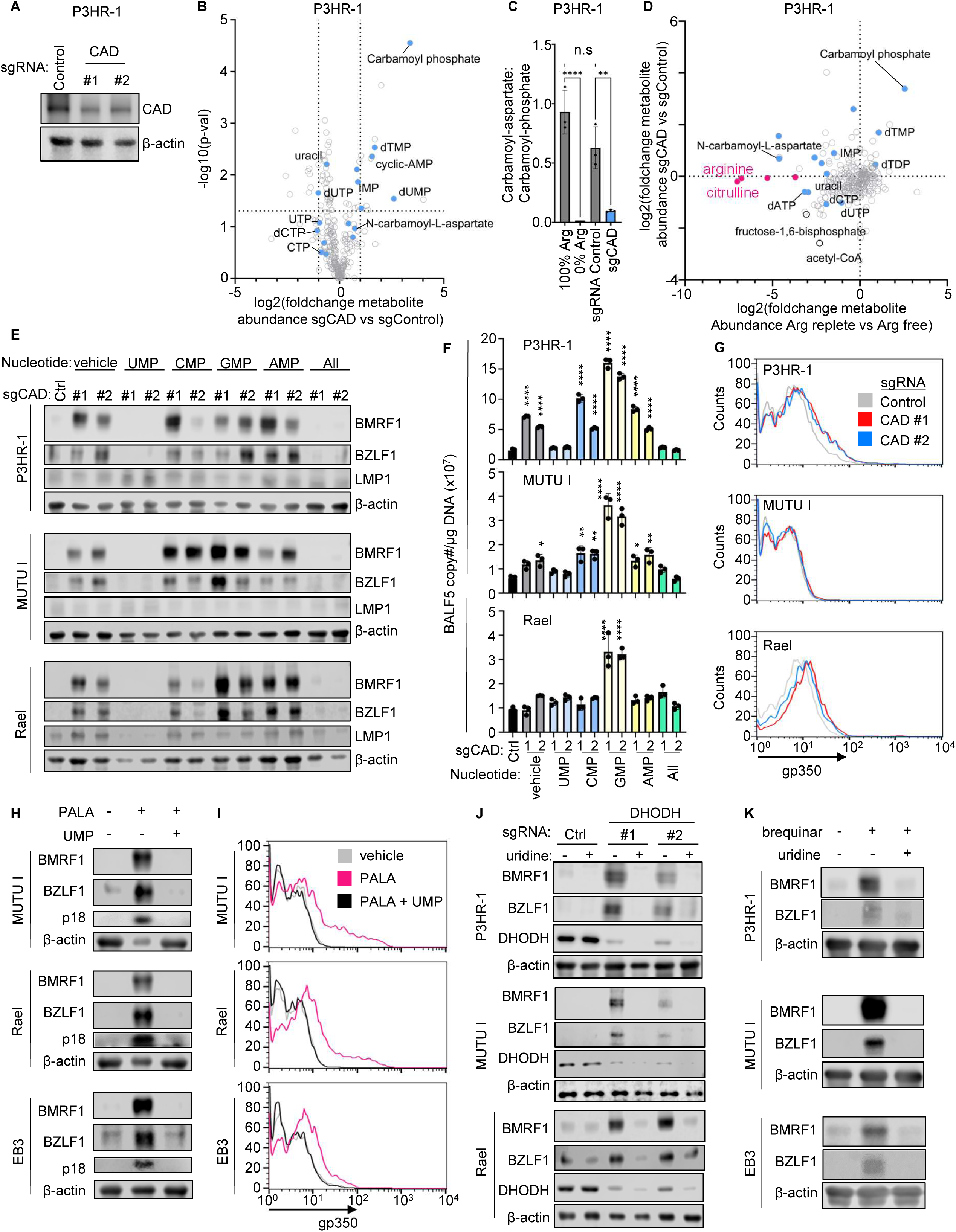
*De novo* pyrimidine biosynthesis is necessary for maintenance of Burkitt EBV latency. (A) Immunoblot analysis of WCL from P3HR-1 cells expressing the indicated control or CAD targeting sgRNAs for 6 days. (B) Volcano plot of LC/MS metabolomic analysis of P3HR-1 cells expressing control or CAD targeting sgRNAs for 6 days from n=3 replicates. Shown are the metabolite foldchange levels from CAD depleted versus control cells. Higher foldchange indicate increased levels in CAD depleted cells. Nucleotide-related metabolites are highlighted in blue. (C) CAD activity, as judged by the carbamoyl-aspartate/carbamoyl-phosphate ratio, in P3HR-1 cultured in arginine free vs replete RPMI media for 5 days, or expressing CAD vs control sgRNAs. (D) Volcano plot of log2 fold-change metabolite abundance from metabolomic analysis of P3HR-1 cells that were cultured in arginine free vs replete media as in (B) (x axis) versus in CAD depleted vs control P3HR-1 (y axis). Arginine cycle metabolites are highlighted in pink; pyrimidine and purine related metabolites are highlighted in blue. (E) Analysis of nucleotide rescue effects on EBV latency in CAD depleted cells. Immunoblot analysis of WCL from cells that expressed control or *CAD* targeting sgRNA and that were cultured in media supplemented with the indicated nucleotides at 50μg/ml, added from 48 hours after sgRNA expression. (F) Analysis of nucleotide rescue effects on EBV lytic genome replication in CAD depleted cells. Shown are mean ± SD values from qPCR analysis of intracellular EBV genome copy number from cells as in (E). (G) FACS analysis of plasma membrane gp350 level in the indicated Burkitt cells that expressed control or *CAD* targeting sgRNA. (H) Immunoblot analysis of WCL from Burkitt cells cultured in RPMI media containing the CAD inhibitor PALA (250μM) and/or UMP (50μg/ml) as indicated for 4 days. (I) FACS analysis of plasma membrane gp350 expression in cells cultured as in (H). (J) Immunoblot analysis of WCL from cells that expressed control or *DHODH* targeting sgRNA and cultured in RPMI supplemented with 50 μg/ml uridine, as indicated. (K) Immunoblot analysis of WCL from cells cultured with the DHODH inhibitor brequinar (BQR, 3μM) and with uridine (50 μg/ml) for 4 days, as indicated. Two-way ANOVA, one-way ANOVA, or Student’s T test was performed, with ****p < 0.0001, ***p < 0.001, **p < 0.01, *p < 0.05. Blots are representative of n=3 replicates.

We next tested whether *de novo* synthesis of pyrimidine biosynthesis is necessary for EBV latency. In support of this hypothesis, CAD KO de-repressed BZLF1 and BMRF1 expressions, which were fully suppressed by UMP supplementation (**Figure 4E**). Importantly, UMP can be metabolized to all other pyrimidines. By contrast, supplementation with the nucleotides cytidine monophosphate (CMP), guanosine monophosphate (GMP) or adenosine monophosphate (AMP) failed to suppress arginine-restriction driven EBV reactivation (**Figure 4E**). Interestingly, CAD KO increased intracellular EBV genome copy number, and this could be suppressed by UMP, but not by CMP, GMP or AMP supplementation (**Figure 4F**). We also noted that GMP supplementation significantly boosted EBV genome copy number in CAD KO cells (**Figure 4F**). CAD KO also modestly increased gp350 expression in P3HR-1 and Rael cells (**Figure 4G**). However, GMP supplementation alone was sufficient to increase EBV genome copy number in control Burkitt cells grown in arginine replete conditions, but not to de-repress EBV lytic antigens, suggesting an effect at the level of latent genome replication (**Figure S4D-E**). Taken together, these results highlight crosstalk between EBV genome replication and nucleotide metabolism pathways.

We next treated Burkitt cells with the well characterized CAD aspartate transcarbamylase inhibitor PALA (N-phosphonacetyl-L-aspartate) (75) to block the first three steps of *de novo* pyrimidine biosynthesis, which produces dihydroorotate from glutamine and aspartate. PALA robustly induced EBV early lytic antigen expression, validating our CRISPR analyses, and in contrast to arginine restriction, also increased late lytic p18 capsid and gp350 protein expression in multiple Burkitt cell contexts. These were on-target effects, because each were rescuable by UMP supplementation (**Figure 4H-I** and **S3F-G**), and even low doses of PALA depresessed BZLF1, BMRF1 LMP1 (**Figure S3H**). Despite UMP rescue of PALA effects on EBV reactivation, UMP supplementation did not maintain EBV latency in arginine restricted cells (**Figure S4I**), possibly indicating additional obligatory arginine metabolism roles in Burkitt EBV maintenance of latency. Alternatively, this could be due to arginine roles in uridine uptake and/or uridine salvage pathway metabolism (76).

We next tested whether the mitochondrial enzyme DHODH, which converts dihydroorotate and quinone to orotate and reduced quinone, and which can form a “pyrimidinosome” complex together with CAD and UMPS across mitochondrial membranes (56), was necessary for EBV latency. We perturbed Burkitt DHODH activity by CRISPR editing or by the small molecule DHODH inhibitor brequinar (77, 78). DHODH inhibition de-repressed BZLF1 and BMRF1, which could be rescued by uridine supplementation, suggestive of on target effects on the *de novo* pyrimidine biosynthesis pathway (**Figure 4J-K**). These results suggest that DHODH orotate synthetase activity, and not quinone reductase activity, was critical for EBV latency.

### Arginine restriction triggers DNA damage and mitochondrial ROS production to reactivate EBV

Given the above observations, we hypothesized that arginine restriction triggers EBV reactivation through effects on nucleotide pool imbalance and DNA damage, the latter of which triggers autophosphorylation by the kinases ATM (ataxia-telangiectasia mutated) and ATR (ATM- and Rad3-Related). ATM is activated by double strand DNA breaks, whereas ATR is activated by single stranded DNA damage or replication stress (79–81). Furthermore, ATM and ATR driven p53 upregulation can reactivate EBV within latently infected gastric cancer cells (82). In support, we found that arginine restriction induced both ATM and ATR phosphorylation across multiple Burkitt cell contexts (**Figure 5A**). Since BZLF1 expression alone can induce DNA damage (83), we also examined DNA damage responses to arginine restriction in EBV-negative Burkitt cells. Interestingly, arginine restriction induced phosphorylation of ATR and its downstream kinase substrate CHK1, but not appreciably of ATM or its substrate CHK2 in EBV-MUTU cells (**Figure 5B**), suggesting a potentially EBV-specific effect on the ATM pathway. To test whether DNA damage responses were required for arginine restriction-induced EBV reactivation, we evaluated effects of ATM inhibitor VE-822 or ATR inhibitor KU-60019. ATR but not ATM inhibition partially suppressed arginine restriction-driven BZLF1 and BMRF1 expression in P3HR-1 (**Figure 5C**). ATM or ATR inhibition partially blocked arginine restriction-driven BZLF1 and BMRF1 but not LMP1 protein induction in EB3 Burkitt cells (**Figure 5C**). These results suggest that the ATM and ATR pathways may each serve as drivers of EBV reactivation downstream of arginine restriction to varying degrees in distinct Burkitt contexts, which we note frequently accumulate mutations in DNA damage pathways (84).

**Figure 5.**
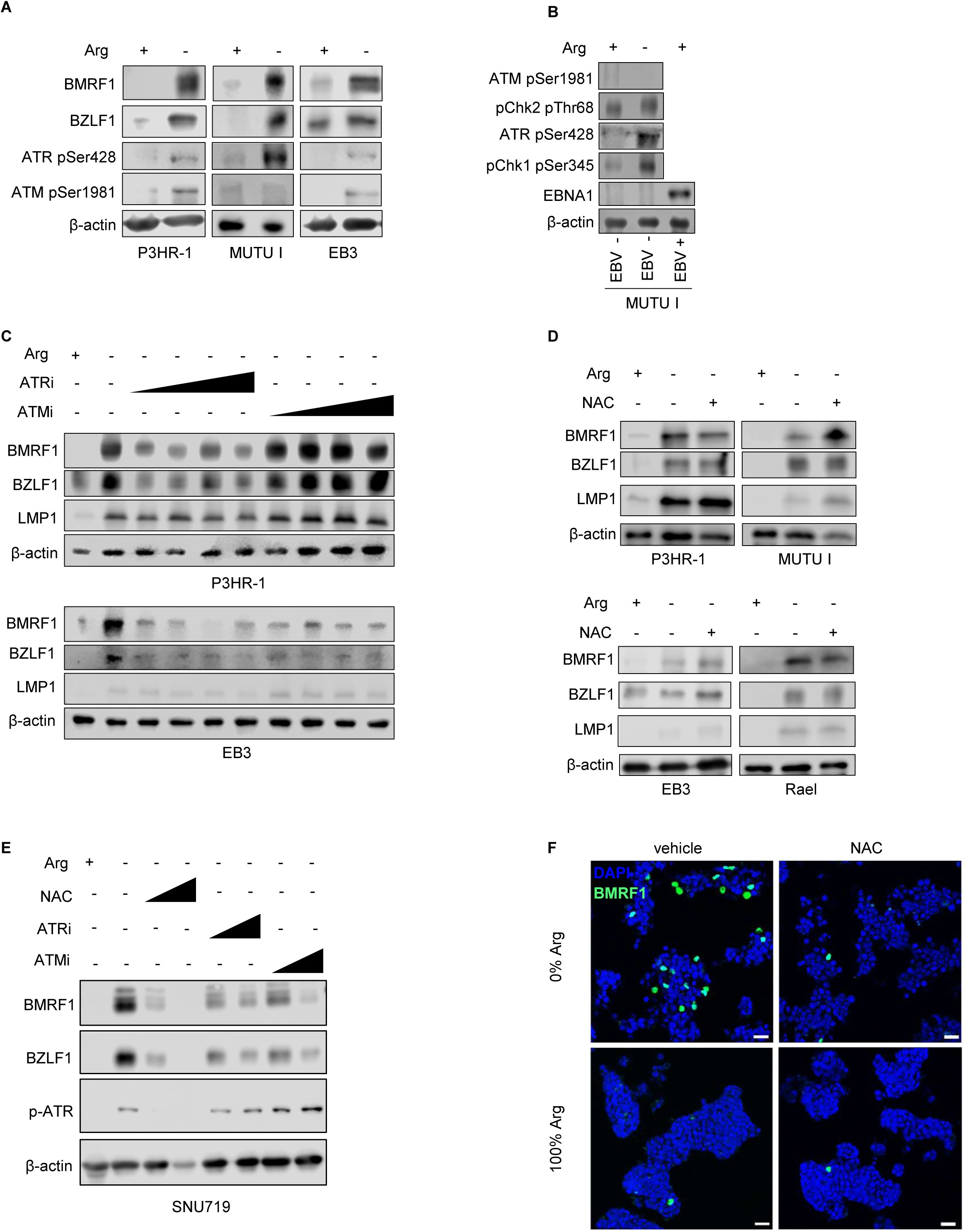
DNA damage responses contribute to EBV arginine restriction driven EBV reactivation. (A) Analysis of arginine restriction effects on ATM and ATR phosphorylation as a readout of their activity. Immunoblot analysis of WCL from P3HR-1, MUTU I or EB3 cells cultured in arginine replete or free media for 5 days. (B) Analysis of arginine restriction effects on DNA damage signaling in EBV-negative Burkitt cells. Immunoblot analysis of WCL from EBV-negative versus positive MUTU I cells cultured for 5 days in arginine replete versus free media. Shown at right as a positive control are EBV+ MUTU I cells. (C) Effects of ATR or ATM inhibition on arginine restriction driven EBV reactivation. Immunoblot analysis of WCL from P3HR-1 or EB3 cells cultured in arginine replete or free media, without or with increasing concentrations of the ATR inhibitor VE-822 (ATRi) or the ATM inhibitor KU-60019 (ATMi) for 5 days. VE-822 and KU-60019 were used at 1.2, 2.5, 5.0 or 10μM. (D) Immunoblot analysis of WCL from P3HR-1 or EB3 cells cultured in arginine replete vs free media with or without 10mM N-acetyl cysteine (NAC) for 5 days. (E) Immunoblot analysis of WCL from SNU-719 gastric carcinoma cells cultured in arginine replete vs free media, without or with increasing concentrations of NAC, VE-822 (5.0μM) or KU-60019 (10μM) for 5 days. NAC was used at 10mM or 50mM final concentration. (F) Representative confocal immunofluorescence images of BMRF1 expression (green) versus nuclear DAPI stain in SNU-719 cells grown in arginine replete or free media for 5 days, in the absence or presence of 10mM NAC as in (E). Scale bars = 50 μm. Blots are representative of n=2 or 3 replicates.

Disruption of mitochondrial membrane potential also inhibits *de novo* pyrimidine biosynthesis at the level of DHODH and activates downstream p53 signaling in part through alteration of reactive oxygen species (ROS) levels (58). To explore potential ROS roles downstream of arginine restriction, we treated Burkitt or SNU719 gastric carcinoma cells with vehicle control or with the antioxidant NAC (N-acetyl cysteine, 10 mM) for 5 days, together with arginine restriction. NAC partially suppressed EBV lytic protein induction in SNU719, but not appreciably in Burkitt cells (**Figure 5D-E**). These results suggest that arginine restriction driven ROS production may contribute to EBV reactivation in particular latently infected epithelial cell contexts, but is not required in Burkitt cells.

### Arginine restriction and DNA hypomethylation jointly de-repress latency I

Many EBV-infected tumors, including Burkitt lymphoma, exhibit CpG island DNA hypermethylation, and DNA hypomethylation also triggers EBV reactivation (13, 74, 85–92). We therefore examined arginine restriction effects on DNA methylation and found that withdrawal of extracellular arginine rapidly increased global Burkitt DNA methylation levels (**Figure 6A**). Yet, methylated DNA immunoprecipitation-qPCR analysis revealed that arginine restriction had variable effects on EBV genomic promoter methylation levels. In P3HR-1, multiple lytic promoters exhibited decreased methylation following 5 days of culture in arginine-free media, even in the presence of acyclovir to block lytic cycle synthesis of unmethylated EBV genomes, while in other two Burkitt cells, the methylation status of most lytic and latency promoters did not change significantly (**Figure 6B, S4A-B**). These results suggest that arginine metabolism differentially regulates host versus EBV genomic DNA methylation, and raised the question of whether arginine restriction and DNA hypomethylation might synergistically or additively trigger EBV reactivation.

**Figure 6.**
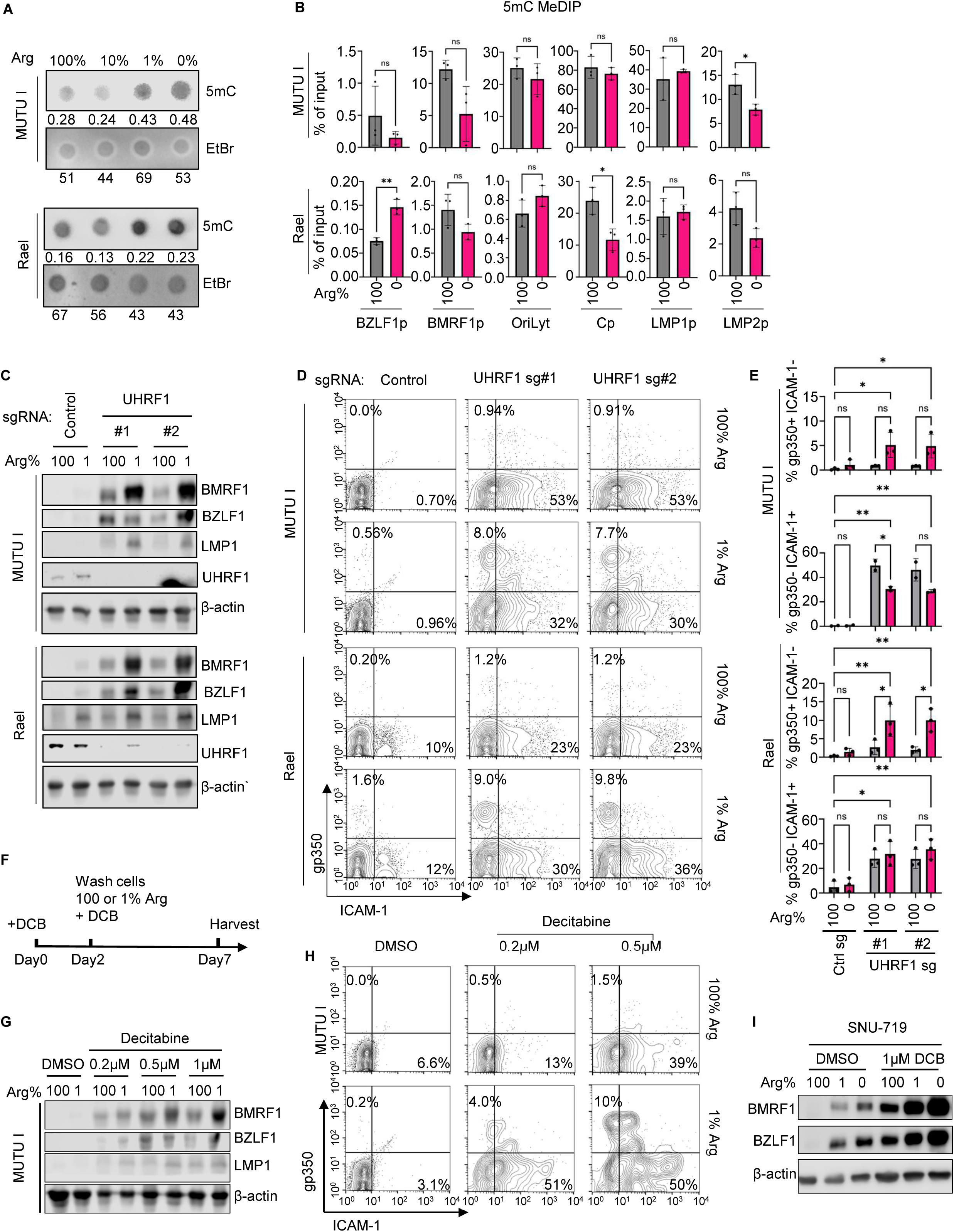
Effects of combinatorial arginine restriction and DNA hypomethylation on EBV reactivation. (A) Analysis of arginine restriction effects on Burkitt 5-methylcytosine (5mC) levels. 5mC dot blot analysis of DNA extracted from MUTU I or Rael cells that were cultured in RPMI with the indicated arginine levels for 5 days. As a load control, membranes were also stained with ethidium bromide (EtBr). Each dot contains 500 ng (MUTU I) or 1 μg (Rael) of DNA. (B) 5mC methylated DNA immunoprecipitation-qPCR (MeDIP) analysis of chromatin from MUTU I or Rael cells cultured in arginine replete vs free media for 5 days. 100μg/ml acyclovir was added to prevent lytic DNA replication. Mean ± SD values from n = 3 replicates are shown. (C) Immunoblot analysis of WCL from MUTU I or Rael cells that expressed control or *UHRF1* targeting sgRNA and cultured in media with 1% or 100% of RPMI arginine levels. Cells were induced to express sgRNA for 3 days and then cultured in indicated media for 5 days. (D) FACS analysis of plasma membrane gp350 expression as a marker of lytic reactivation vs ICAM-1 expression as a marker of LMP1 expression in cells cultured as in (C). (E) Mean ± SEM percentages of gp350+ or ICAM+ cells from n=3 experiments as in (D). (F) Schematic of arginine restriction and decitabine (DCB) treatments. (G) Immunoblot analysis of WCL from MUTU I cells that were cultured in media containing the indicated DCB concentrations for 2 days and then cultured in arginine restricted (1% RPMI levels) or replete media for 5 days. (H) FACS analysis of plasma membrane gp350 or ICAM-1 expression on cells cultured as in (G). (I) Immunoblot analysis of WCL from SNU-719 gastric carcinoma cells that were cultured in media containing the indicated DCB concentrations for 2 days and then cultured in arginine restricted (1% Arg) or replete media for 5 days. Two-way ANOVA or Student’s t test was performed, with ****p < 0.0001, ***p < 0.001, **p < 0.01, *p < 0.05. Blots are representative of n=3 replicates.

CpG DNA methylation marks are propagated onto newly synthesized Burkitt B cell host and EBV genomes by the host cell enzymes DNMT1 and UHRF1 (85). We therefore next tested the effects of UHRF1 depletion on EBV reactivation in P3HR-1, MUTU I, and Rael cells, alone or together with arginine restriction for five days. Interestingly, UHRF1 depletion and arginine restriction more highly induced MUTU I and Rael lytic antigen expression than either did alone, suggestive of additive or synergistic effects on EBV reactivation (**Figure 6C**).

We reported that DNA hypomethylation resulted in distinct populations of Burkitt cells with EBV expression (gp350+) versus with de-repression of the latency III program, in which LMP1 upregulated plasma membrane ICAM-1 expression (85, 93). In MUTU I and Rael cells, UHRF1 knockout and arginine restriction together upregulated the percentage of gp350+/ICAM-1-cells, indicative of reactivation and potentially full lytic replication, given late lytic protein expression (**Figure 6E**). We note that this analysis may have underestimated the percentage of cells with abortive lytic EBV reactivation. UHRF1 KO and arginine restriction also significantly increased the percentage of gp350-/ICAM-1+ cells, suggestive of LMP1 de-repression (**Figure 6E**), potentially with switch to latency III. Synergistic gp350 induction was not observed in P3HR-1 cells, likely because arginine restriction alone strongly hypomethylated viral genome promoters (**Figure S5C-E**). The hypomethylating agent decitabine and arginine restriction also showed synergistic effects on the induction of BZLF1, BMRF1 and even late gp350 expression (**Figure 6G-H, S5F-G**), including in SNU-719 gastric carcinoma cells (**Figure 6I**). Taken together, these data indicate that arginine restriction and DNA hypomethylation each strongly support the maintenance of the latency I program, and that targeting them concomitantly enhances the switch to lytic and latency III programs within distinct single cell populations.

To further test potential therapeutic combinations, we further defined effects of concomitant CAD inhibition and DNA hypomethylation. MUTU I and Rael cells were treated with decitabine for 2 days and then with PALA for 4 days. At the 0.2μM decitabine dose, PALA strongly increased EBV lytic protein expression in MUTU I and Rael (**Figure 7A**). FACS analysis indicated that 0.2 μM decitabine/PALA co-treatment together significantly increased the %gp350+ cells, particularly in Rael cells (**Figure 7B**), again suggesting a full lytic cycle induced by PALA and increased by decitabine co-administration. Taken together, our results support a model in which arginine metabolism supports the maintenance of the EBV latency I state through feeding the *de novo* pyrimidine synthesis pathway, whose production of UMP prevents DNA damage. Interruption of either arginine metabolism or UMP biosynthesis triggers EBV reactivation, more strongly when applied together with DNA hypomethylation (**Figure 7C**).

**Figure 7.**
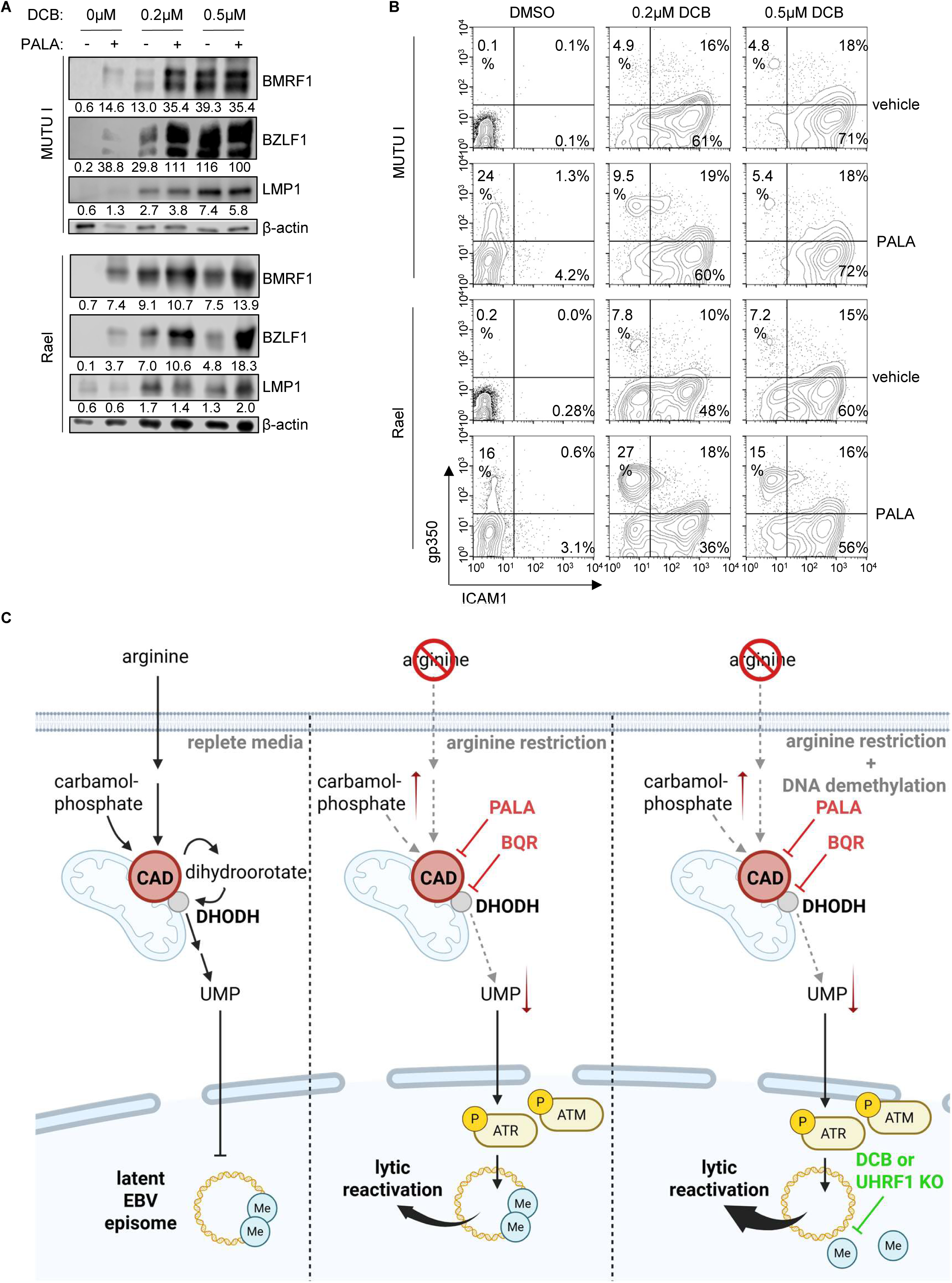
PALA and DCB synergize to induce EBV lytic antigen expression. (A) Immunoblot analysis of WCL from MUTU I or Rael cells that were treated with the indicated DCB concentrations for 2 days and then also with vehicle versus PALA (250μM) for 4 days, as indicated. (B) FACS analysis of cells treated as in (A). (C) Schematic model of arginine and of *de novo* pyrimidine metabolism effects on EBV latency, alone or together with DNA hypomethylation.

## Discussion

EBV senses the infected cell microenvironment and physiological state to dictate whether to remain latent or to reactivate, which enables viral spread but risks immune detection. Although they are a major regulator of cells nutritional status, much has remained unknown about how cross-talk between amino acid metabolic pathways and the EBV epigenome regulates the viral lytic switch. Here, we build on a classic observation by founders of the EBV field to investigate the mechanism by which arginine metabolism supports EBV latency within Burkitt B cells and other EBV-infected cell contexts. Metabolomic profiling highlighted that arginine metabolism supports *de novo pyrimidine* synthesis, which was necessary to support infected cell levels of uridine monophosphate, the key precursor for all pyrimidine nucleotides. Restriction of extracellular arginine or inhibition of the rate-limiting *de novo pyrimidine* biosynthesis enzyme CAD, which is supplied by upstream arginine metabolism, activated ATM and ATR driven DNA damage signaling. Arginine restriction and DNA hypomethylation together more highly de-repressed EBV latency I than either alone, suggesting novel therapeutic approaches.

We note that restriction of either methionine (74) or arginine de-repressed EBV immediate early and early lytic genes, but not late genes, suggesting roles for arginine and methionine metabolism in synthesis of lytic EBV genomes, the DNA templates for late gene expression. Interestingly however, PALA de-repressed expression of the EBV late genes p18/viral capsid antigen and gp350, even though we did not observe late gene expression with CRISPR CAD KO. PALA is a rationally-synthesized analog of the transition-state intermediate produced by CAD conversion of carbamoyl phosphate and aspartic acid into carbamoyl aspartate (94). Although PALA is a potent CAD inhibitor with an estimated Km of ∼2 x 10^-5^ M (95), it is nonetheless competitive with carbamoyl phosphate. We therefore hypothesize that partial CAD inhibition by PALA is sufficient to constrain *de novo* pyrimidine biosynthesis, as evidenced by uridine rescue, but still allows for some enzyme activity needed for lytic viral genome synthesis and late gene expression.

With regards to the mechanism by which inhibition of CAD or *de novo* pyrimidine biosynthesis triggers EBV reactivation, our data suggests that arginine restriction triggers DNA damage signaling. This may arise from imbalanced nucleotide pools and replication stress (96, 97) and/or from increased mitochondrial production of ROS, which can oxidize nucleotides, can directly activate ATM (98) and can cause single strand DNA breaks (81). Amino acid restriction, including that of arginine, inhibits mTOR (99). Of these, our data favor nucleotide imbalance as the key factor at least in Burkitt cells, because arginine restriction activated both ATM and ATR pathways, and arginine restriction effects on lytic gene expression could be partially reversed by ATR or ATM inhibition or by UMP supplementation. UMP, but not other nucleotide monophosphates, rescued EBV latency in arginine restricted cells. UMP can be metabolized to each of the other pyrimidine nucleotides. By contrast, CMP is not readily converted to UMP (100). Therefore, it is likely that arginine and CAD support of multiple pyrimidine nucleotide pools contributes to EBV latency, rather than support of UMP alone.

There is significant interest in EBV lytic induction therapies, and recent clinical trials have shown early but promising results (32, 101). Exploiting EBV-infected cell metabolic vulnerabilities provides an opportunity to sensitize EBV-infected cells to the antiviral ganciclovir, whose cytotoxicity is activated by the EBV early lytic kinase BGLF4 (26). Furthermore, dependence on extracellular arginine is a metabolic vulnerability in many types of cancers, and PEGylated arginase is in clinical development for multiple types of cancers (102). It will therefore be of interest to test the extent to which arginase counteracts growth of tumors in Burkitt or gastric carcinoma xenograft models, either alone or in combination with hypomethylation and/or ganciclovir.

We recently found that restriction of extracellular methionine altered the EBV epigenome, reducing CpG DNA methylation and altering several histone epigenetic marks (74). While arginine restriction likewise de-repressed EBV immediate early and early lytic antigens, we instead found that arginine restriction increased global Burkitt 5-methylcytosine levels, though interestingly had variable effects on EBV genomic DNA methylation. We speculate that this property underlies our observation that arginine restriction and DNA hypomethylation by UHRF1 KO or by decitabine together more highly induced EBV lytic reactivation than either did alone. Interestingly however, as we previously described (74, 93), Burkitt cells in the latency I state appear to have a stochastic choice of switching to lytic reactivation versus to latency III. Yet, whereas decitabine or UHRF1 KO trigger a Burkitt subpopulation to de-repress LMP1 and its target ICAM-1, PALA did not induce ICAM-1 expression and instead preferentially de-repressed gp350, indicative of lytic reactivation. We note that plasma membrane gp350 staining likely underestimated the extent of lytic reactivation, given that cells with an abortive lytic cycle may have expressed immediate early and early EBV lytic proteins but not late proteins. Since PALA and decitabine induced most cells to either derepress the lytic cycle or LMP1, an intriguing approach would be to follow PALA/decitabine treatment with ganciclovir and to then adoptively transfer LMP1-specific cytotoxic T-cells (CTL). This might serve to improve responses beyond those observed with decitabine and CTL therapy alone, where T-cells homed to low dose decitabine-treated tumors and exerted a degree of anti-tumor effect at 3 weeks post-infusion (93).

In summary, EBV-infected tumor cells rely on arginine metabolism to support the maintenance of EBV latency in both B and epithelial cell contexts. Mechanistically, extracellular arginine supports CAD-driven *de novo* pyrimidine synthesis to supply sufficient UMP, which can then be converted to all of the pyrimidine nucleotides. Arginine restriction therefore caused nucleotide pool imbalance and DNA damage, which in turn triggered reactivation. Arginine restriction or CAD inhibition and DNA hypomethylation together induced higher levels of EBV reactivation, suggesting therapeutic approaches.

## Acknowledgements

This work was supported by T32 AI007061, by American Cancer Society Post-doctoral Fellowships PF-24-1308318-01-TBE to S.W and PF-23-898493-01-TBE to E.M.B, by Lymphoma Research Foundation Postdoctoral Fellowship to Y.L., and by R01AI164709, R01DE033907, R01CA228700, U01CA275301 and P01CA269043 to B.E.G.

## AUTHOR CONTRIBUTIONS

S.W performed the experiments. S.W, Y.L and E.M.B performed bioinformatics analyses. S.W and B.E.G designed the experiments. S.W and B.E.G wrote the manuscript. All authors analyzed the results, read and approved the manuscript.

---

**Figure S1. Arginine or histidine restriction effects on EBV+ Burkitt cells, related to Figure 1**.

(A) Effects of essential amino acid (AA) restriction on P3RH-1 proliferation. Shown are mean ± standard error of the mean (SEM) fold-change in live cell number of cells grown in replete media versus in media lacking the indicated essential amino acid, normalized to cell numbers at day 1. (B) Effects of non-essential AA restriction on P3RH-1 proliferation. Although considered non-essential amino acids that do not have be acquired by diet, restriction of either cystine, arginine, glutamine or tyrosine significantly impaired P3HR-1 proliferation. (C) Immunoblot analysis of immediate early BZLF1, early BMRF1, phospho-p70 S6 kinase (Thr389), total p70S6 kinase or load control β-actin using WCL from P3HR-1 cells cultured in media with vehicle or 10 nM rapamycin for 3 or 5 days. (D) Immunoblot analysis of WCL from P3HR-1 cells cultured in replete or arginine-free media for 5 days, with or without 100 μg/ml acyclovir (ACV) to block EBV lytic replication. (E) Histidine restriction effects on EBV lytic reactivation. Immunoblot analysis of WCL from P3HR-1, MUTU I or EB3 Burkitt cells cultured in media for three days with the indicated histidine level, where 100% histidine refers to the typical RPMI histidine concentration (97μM).

**Figure S2. Analysis of arginine restriction effects on P3HR-1 Burkitt growth and survival.**

(A) Analysis of arginine restriction effects on P3HR-1 Burkitt B cell proliferation. Growth curves of P3HR-1 cells cultured in media with the indicated arginine (Arg) concentration. 100% arginine refers to the RPMI arginine concentration of 115 μM. Mean ± SD values from n = 4 replicates are shown. (B) FACS analysis of cells in (A).

**Figure S3. Arginine restriction effects on ATP, ADP and AMP, related to Figure 3**.

(A) Analysis of arginine restriction effects on adenosine nucleotide levels, from LC/MS metabolomic analysis of EB3 cells grown in replete (100% Arg) vs arginine free (0% Arg) media, as in Figure 3A. (B) Analysis of arginine restriction effects on the ATP/ADP or ADP/AMP ratios, from the LC/MS analysis presented in Figure 3A. (C) Analysis of arginine restriction effects on ASS1 expression. Immunoblot analysis of WCL from the indicated Burkitt cells cultured in arginine replete vs free RPMI for 5 days.

**Figure S4. *De novo* pyrimidine biosynthesis maintains EBV latency, related to Figure 4**.

(A) Metabolic analysis of P3HR-1 cells cultured in arginine-free vs replete RPMI for 5 days. Shown are foldchanges of metabolite abundance and p-values, calculated from n=3 replicates. Arginine related metabolites are highlighted in pink; pyrimidine and purine related metabolites are highlighted in blue. Higher foldchange indicate higher metabolite abundance in cells grown in arginine replete than arginine free media. (B) KEGG metabolic pathway analysis of metabolomic data as in (A). Metabolites were selected using a FDR<0.05 cutoff and pathway impact values were computed by MetaboAnalyst 3.0 topological analysis. (C) Volcano plot visualization of log2 foldchange in metabolite abundance from LC/MS analysis of P3HR-1 cells (x-axis) versus of EB3 cells (y-axis) cultured in arginine replete vs free media for 5 days. Arginine cycle metabolites are highlighted in pink; pyrimidine and purine related metabolites are highlighted in blue. (D) Immunoblot analysis of WCL from P3HR-1 cells that were cultured in RPMI supplemented with the indicated nucleotide monophosphates to a total concentration of 50μg/ml. Shown at right as a + control or lysates from CAD depleted cells, taken at 48 hours after CAD-targeting sgRNA expression. (E) qPCR analysis of intracellular EBV genome copy number of cells as in (D). Shown are the mean ± SD values from n=3 replicates. (F) Immunoblot analysis of WCL from P3HR-1 cells cultured in media with 250μM PALA and with vehicle versus 50μg/ml UMP for 4 days, as indicated. (G) FACS analysis of plasma membrane gp350 abundance in cells as in (F). (H) Immunoblot analysis of WCL from MUTU I or Rael cells cultured in arginine replete or free media, in the absence or presence of 250μM PALA for 4 days. (I) Immunoblot analysis of WCL from cells cultured in arginine replete or arginine free media with 0μg/ml, 0.5μg/ml, 5μg/ml, 50μg/ml or 500μg/ml UMP supplementation for 5 days. Blots are representative of n=3 replicates.

**Figure S5. Arginine restriction and DNA hypomethylation combinatorial effects on EBV reactivation, related to Figure 5**.

(A) 5mC dot blot analysis of DNA extracted from P3HR-1 cells that were cultured in RPMI with the indicated arginine levels for 5 days. Membranes were stained with ethidium bromide as load controls. Each dot contains 500 ng DNA. (B) 5mC MeDIP analysis of chromatin from P3HR-1 cells cultured in arginine free versus replete RPMI media for 5 days. 100 μg/ml acyclovir was added to prevent lytic DNA replication. Mean ± SD values from n = 3 replicates are shown. (C) Immunoblot analysis of WCL from P3HR-1 cells that expressed control or *UHRF1*-targeting sgRNA and were cultured in arginine restricted (1% Arg) vs replete media. Cells were induced to express sgRNA for 3 days and then cultured in the indicated media for 5 days. (D) FACS analysis of plasma membrane gp350 and ICAM-1 expression on cells treated as in (C). (E) Mean ± SEM percentages of gp350+ and ICAM-1+ cells from n=3 replicates from n=2 independent experiments. (F) Immunoblot analysis of WCL from Rael or EB3 cells that were cultured in media containing indicated DCB concentrations and then cultured in 1% Arg or 100% Arg media for 5 days. (G) FACS analysis of plasma membrane gp350 and ICAM-1 levels in Rael cells treated as in (F). Two-way ANOVA or Student’s T test was performed, with ****p < 0.0001, ***p < 0.001, **p < 0.01, *p < 0.05.

## Materials and Methods

### Cell lines, culture, and growth curve analysis

293T cells were purchased from ATCC and cultured in Dulbecco modified Eagle medium (Gibco) with 10% FBS (Thermo Fisher). Burkitt lymphoma cell lines P3HR-1, EBV-Akata, EBV-MUTU, MUTU I, Rael, EB3, and EBV+ gastric cancer cell line SNU719 were cultured in RMPI 1640 (Gibco, Life Technologies) supplemented with 10% fetal bovine serum (Gibco). P3HR-1 that can be induced to lytic reactivation with 4-HT contains ZHT/RHT, which are BZLF1/BRLF1 fused to a modified estrogen receptor 4HT-binding domain (Gift from Eric Johannsen). P3HR-1, MUTU I, and Rael expressing *Streptococcus pyogenes* Cas9 genes were generated previously by lentiviral transduction, followed by blasticidin selection at 5 μg/ml (103). Cells were tested routinely to be mycoplasma free with the MycoAlert kit (Lonza).

For the single amino acid restriction screen, P3HR-1 cells were cultured in RPMI 1640 medium without amino acids (Thermo Fisher) that is supplied with 10% dialyzed FBS, and supplied with a panel of amino acids to normal RPMI concentrations, excluding the restricted amino acid. For arginine restrictions, cells were cultured in RPMI 1640 Medium for SILAC (Thermo Fisher) with supplementation of normal media concentration (0.22 mM) of L-Lysine and 10% dialyzed FBS, as RPMI 1640 Medium for SILAC doesn’t contain L-arginine and L-lysine. Catalog no. of chemicals can be found in Table S3. For cell growth curve, 10 ml culture in T25 flasks was mixed by pipetting, from which 1 ml culture was taken, centrifuged at 1,500 rpm for 5min, and resuspended in 1ml phosphate-buffered saline (PBS). Cells were stained with trypan blue (Thermo Fisher) and live cell numbers were counted by light microscope.

### Immunoblot analysis

Whole cell lysate (WCL) samples were separated by SDS-PAGE, transferred onto nitrocellulose filters (Bio-Rad), blocked with TBST (Tris-buffered saline with Tween 20) containing 5% milk or 3% BSA (NEB), and then probed with primary antibodies at 4°C overnight, followed by secondary antibody (Cell Signaling Technology, catalog no. 7074, catalog no. 7076, catalog no. 7077; LI-COR INC, catalog no. 926-32210, catalog no. 926-32219) incubation for 1 h at room temperature. For all horseradish peroxidase (HRP)-conjugated secondary antibodies, ECL chemiluminescence (Millipore, catalog no. WBLUF0500) was used to develop HRP signal. All images were captured with a Li-Cor Fc platform. All antibodies used in this study are listed in Table S3.

For DNA dot blot, DNA harvested by DNeasy Blood & Tissue Kit (Qiagen) from cells was hybridized on the nitrocellulose membrane and blocked with TBST containing 5% milk. The membrane was then washed and blotted with anti-5 methyl-cytosine monoclonal antibody overnight or stained with 1 mg/ml ethidium bromide in PBS for 10 min. After further washing with TBST or PBS, membranes were imaged as described above. Densitometry of Western blots were analyzed with Image Studio Ver 5.5. Equal area was selected for each band of the same protein, and relative protein abundances were standardized to corresponding β-actin (Biolegend, 664802) bands for SDS-PAGE blot or to ethidium bromide dot for dot blot.

### Quantification of EBV genome copy number by PCR analysis

Total DNA from 0.5-1×10^6^ cells were extracted with DNeasy Blood& Tissue Kit (Qiagen). Then, the DNA extracted from each sample was diluted to 10 ng/ml before subjected to qPCR analysis with a primer pair targeting the EBV BALF5 gene (Table S3). The quantitative real-time PCR was then performed using the Power SYBR Green PCR Master Mix (Applied Biosystems) on a CFX96 Touch Real-Time PCR Detection System (BioRad). Viral DNA copy number was calculated by interpolation of Cq values to a standard curve generated by serial dilutions of 25 ng/ml pHAGE-BALF5 plasmid.

### Immunofluorescent confocal imaging

5×10^5^ cells suspended in PBS were allowed to dry on glass slides and then fixed with 4% paraformaldehyde (PFA) in PBS for 10 min. Cells were then washed and permeabilized with 0.1% Triton-X in PBS for 5 min. Cells were then blocked with PBS containing 1% low IgG BSA (MPbio) overnight at 4°C. Cells were incubated with primary antibodies for 1 hr at 37°C, washed three times, and incubated in secondary fluorescent antibody for 1hr at 37°C before washed three times again and mounted with ProLong™ Gold anti-fade mountant with DAPI (Thermo Fisher). Confocal images were taken on a Zeiss LSM 800 instrument. BMRF1+ cells were counted by eye with the assistance of FIJI ImageJ software.

### Flow cytometry analysis

All cells were washed once with PBS before staining. For 7-AAD and annexin V flow cytometry analysis, 1×10^6^ cells were stained in 100 μl of staining buffer that contained FITC-conjugated anti-annexin V antibody (1:20) and 5 μg 7-AAD for 15 min. Annexin V antibody and 7-AAD were diluted in annexin V binding buffer, which contains 10 mM HEPES pH 7.4, 140 mM NaCl, and 2.5 mM CaCl_2_. Before flow cytometry analysis, the total volume was expanded to 500 μl with annexin V binding buffer. For gp350 and ICAM1 flow cytometry analysis, cells were incubated with Cy5-conjugaged anti-gp350 antibody (1:800) and PE-conjugated anti-CD54 antibody (1:800) in PBS containing 2% FBS for 45 mins at 4°C. Cells were then washed twice with PBS containing 2% FBS and resuspended in 500 μl PBS containing 2% FBS before flow cytometry analysis. Flow cytometry of cells was performed on a BD FACSCalibur instrument and the analysis was performed with FlowJo V10 software.

### Cellular metabolomic analysis

For profiling the metabolome of arginine restricted EB3 cells, 1×10^7^ cells were seeded into each T75 flask (five flasks for each condition) with 20 ml RPMI 1640 Medium for SILAC, supplied with 10% dialyzed FBS and 0.22 mM L-Lysine. L-arginine was added to 1.15 mM for control (100% Arg) group. Three days after seeding, media was refreshed for all groups by pelleting cells at 1,500 rpm for 10 min and resuspension in equal volume fresh respective media. Cells in control group were split 1:5 to account for cell proliferation. Five days after seeding, 3×10^6^ total cells from each T75 flask were seeded into a T25 in 10 ml respective media 2 hr prior to intracellular metabolite extraction. For profiling the metabolome of arginine restricted P3HR-1 cells, cells were treated the same as EB3 cells, except with three replicates.

For profiling the metabolome of arginine restricted EB3 cells, 1×10^7^ cells were seeded into each T75 flask (five flasks for each condition) with 20 ml RPMI 1640 Medium for SILAC, supplied with 10% dialyzed FBS and 0.22 mM L-Lysine. L-arginine was added to 1.15 mM for control (100% Arg) group. Three days after seeding, media was refreshed for all groups by pelleting cells at 1,500 rpm for 10 min and resuspension in equal volume fresh respective media. Cells in control group were split 1:5 to account for cell proliferation. Five days after seeding, 3×10^6^ total cells from each T75 flask were seeded into a T25 in 10 ml respective media 2 hr prior to intracellular metabolite extraction. For profiling the metabolome of arginine restricted P3HR-1 cells, cells were treated the same as EB3 cells, except with three replicates. For profiling the metabolome of CAD-knockout P3HR1, 3×10^6^ control or CAD depleted cells were collected nine days after CRIPSR editing. Cells were seeded in 10 ml fresh RPMI 1640 media supplied with 10% FBS 2 hr prior to intracellular metabolite extraction.

For metabolite extraction, cells were washed with 5 mL of room temperature PBS. Then, pellets were resuspended in 1 mL of dry ice cold 80% methanol, incubated at −80°C for 30 min and centrifuged at 21,000 x g for 5 minutes at 4°C. Supernatants were collected in pre-chilled tubes and stored at −80°C. 6 replicates were included for each treatment. On the day of analysis, supernatants were incubated on ice for 20 min, clarified by centrifugation at 21,000 x g at 4°C, and dried down with a speed vacuum concentrator (Savant SPD 1010, Thermofisher Scientific). Samples were re-suspended in 100μL of 60/40 acetonitrile/water, vortexed, sonicated in ice-cold water, and incubated on ice for 20 min. Following centrifugation at 21,000xg for 20 min at 4°C, supernatants were collected for pooled QC. Metabolite profiling was performed at the Beth Israel Deaconess Mass Spectrometry Core. Samples were re-suspended using 20 uL HPLC grade water for mass spectrometry. 5-7 μL were injected and analyzed using a hybrid 6500 QTRAP triple quadrupole mass spectrometer (AB/SCIEX) coupled to a Prominence UFLC HPLC system (Shimadzu) via selected reaction monitoring (SRM) of a total of 300 endogenous water soluble metabolites for steady-state analyses of samples. Some metabolites were targeted in both positive and negative ion mode for a total of 311 SRM transitions using positive/negative ion polarity switching. ESI voltage was +4950V in positive ion mode and – 4500V in negative ion mode. The dwell time was 3 ms per SRM transition and the total cycle time was 1.55 seconds. Approximately 9-12 data points were acquired per detected metabolite. Samples were delivered to the mass spectrometer via hydrophilic interaction chromatography (HILIC) using a 4.6 mm i.d x 10 cm Amide XBridge column (Waters) at 400 uL/min. Gradients were run starting from 85% buffer B (HPLC grade acetonitrile) to 42% B from 0-5 minutes; 42% B to 0% B from 5-16 minutes; 0% B was held from 16-24 minutes; 0% B to 85% B from 24-25 minutes; 85% B was held for 7 minutes to re-equilibrate the column. Buffer A was comprised of 20 mM ammonium hydroxide/20 mM ammonium acetate (pH=9.0) in 95:5 water:acetonitrile. Peak areas from the total ion current for each metabolite SRM transition were integrated using MultiQuant v3.0.2 software (AB/SCIEX). Metabolites with CV<30% in pooled QC were used for the statistical analysis. The quality of integration for each metabolite peak was reviewed. Metabolites with p-values < 0.05, log2(fold change)>1 or <-1 were used for pathway analysis using MetaboAnalyst 5.0 (https://www.metaboanalyst.ca/MetaboAnalyst/ModuleView.xhtml). Heatmaps of metabolites in the pathways were generated by feeding Z-score values into Morpheus software (https://software.broadinstitute.org/morpheus/).

A separate cell pellet from each intracellular metabolite extraction sample was then quantified for protein content using a BCA assay kit (Thermofisher Scientific). Protein content was calculated by interpolation of OD_562_ values to a standard curve generated by serial dilutions of bovine serum albumin provided in the kit. Protein content ratios between samples were calculated by normalization, which was used to guide the standardization of metabolites peaks. Metabolites with p-values < 0.05, log_2_ fold-change >1 or <-1 were used for pathway analysis using MetaboAnalyst 6.0.

### CRISPR/Cas9 Mutagenesis

CRISPR/Cas9 editing was performed as previously described (103). Briefly, Broad Institute pXPR-011 GFP-targeting, sgRNA non-targeting control (BRDN0004438510), Avana or Brunello library sgRNAs were cloned into shuttle vector lentiGuide-Puro (Addgene, #52963) or pRDA_355 (Addgene, #187159). Production of lentivirus and transduction of target cells for CRISPR/Cas9 gene editing was described before (104). Shuttle vector, packaging vectors psPAX2 and VSV-G were transfected into 293T cells with TransIT-LT1 Transfection Reagent (Mirus) according to the manufacturer’s protocol. 12 hr post transfection, cell culture media was changed to RPMI-1640 with 10% FBS. 293T supernatants were harvested 48 and 72 hr post transfection, and were dripped into target cells that were seeded at 3×10^5^ cells/ml. Cells were selected by puromycin (3 mg/mL), added 48 h post-transduction. For CRISPR/Cas9 knockout using pRDA_355, doxycycline (Sigma) was added to 200 ng/ml to induce the expression of sgRNA. Cell pellets were harvested 48 hr post CRISPR editing and analyzed with immunoblot as described above to confirm the loss of protein expression.

### 5-methyl cytosine DNA immunoprecipitation (MeDIP) and qPCR

Genomic DNA was purified using the Blood and Cell Culture DNA Mini Kit (Qiagen) and then used for MeDIP analysis, using the MagMeDIP kit (Diagenode cat #C02010021), according to the manufacturer’s protocol. qPCR assays were then performed as described above.

### Software/data Presentation

Graphs were made using GraphPad Prism 10. Schematic models were made using Biorender.

