## supplemental figures and tables for "Arginine Metabolism Supports *De Novo* Pyrimidine Biosynthesis to Block DNA Damage and Maintain Epstein-Barr Virus Latency": Arginien_restriction_sup_figures.pdf

**Figure S1**

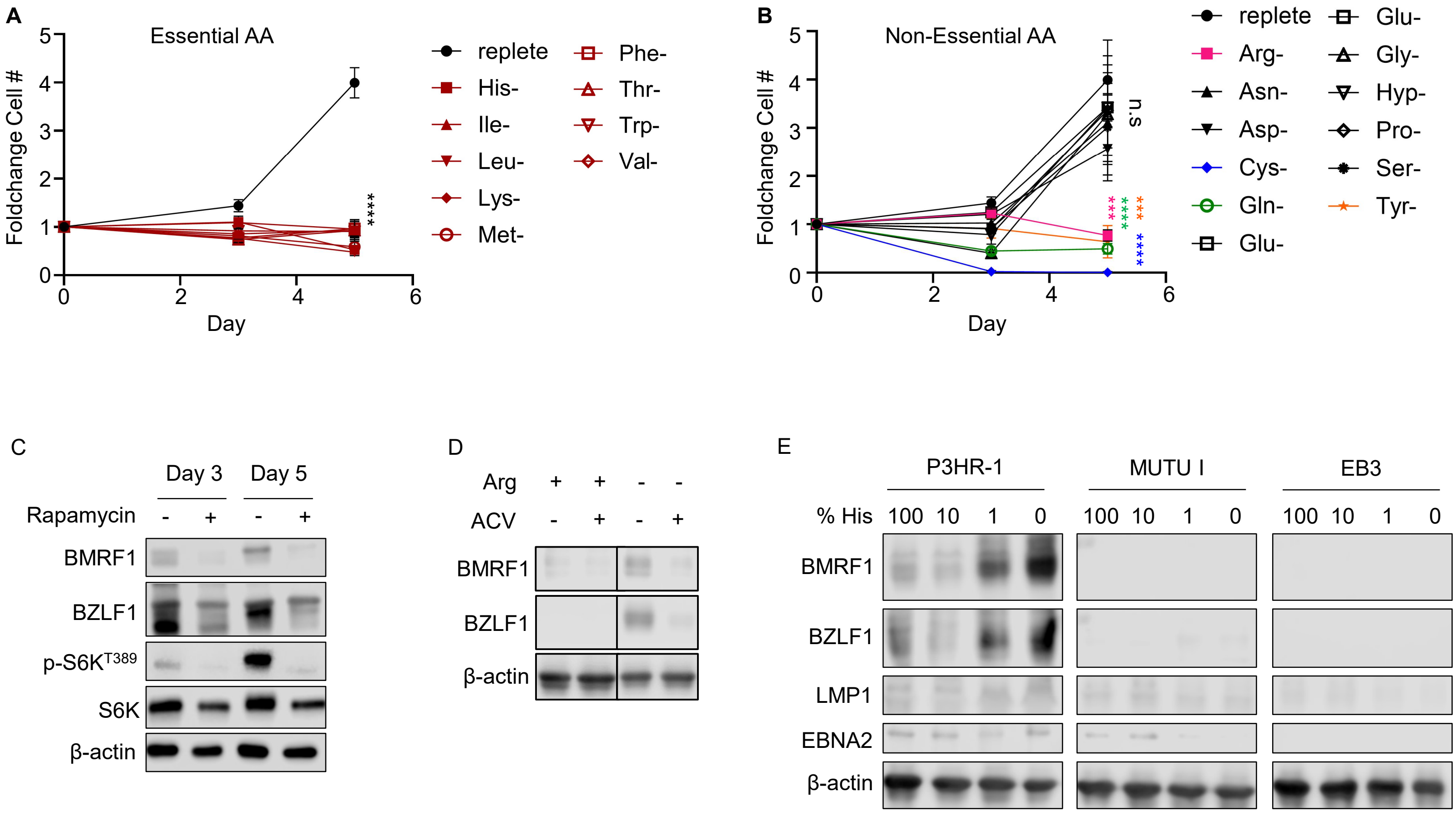
A
Essential AA
Foldchange Cell #
Day
****
replete
His-
Ile-
Leu-
Lys-
Met-
Phe-
Thr-
Trp-
Val-
B
Non-Essential AA
Foldchange Cell #
Day
n.s
***
****
***
****
replete
Arg-
Asn-
Asp-
Cys-
Gln-
Glu-
Glu-
Gly-
Hyp-
Pro-
Ser-
Tyr-
C
Day 3
Day 5
Rapamycin
BMRF1
BZLF1
p-S6K T389
S6K
β-actin
D
Arg
ACV
BMRF1
BZLF1
β-actin
E
P3HR-1
MUTU I
EB3
% His
100 10 1 0
BMRF1
BZLF1
LMP1
EBNA2
β-actin

**Figure S1. Arginine or histidine restriction effects on EBV+ Burkitt cells, related to Figure 1.**
(A) Effects of essential amino acid (AA) restriction on P3RH-1 proliferation. Shown are mean ± standard error of the mean (SEM) fold-change in live cell number of cells grown in replete media versus in media lacking the indicated essential amino acid, normalized to cell numbers at day 1. (B) Effects of non-essential AA restriction on P3RH-1 proliferation. Although considered non-essential amino acids that do not have be acquired by diet, restriction of either cystine, arginine, glutamine or tyrosine significantly impaired P3HR-1 proliferation. (C) Immunoblot analysis of immediate early BZLF1, early BMRF1, phospho-p70 S6 kinase (Thr389), total p70S6 kinase or load control β-actin using WCL from P3HR-1 cells cultured in media with vehicle or 10 nM rapamycin for 3 or 5 days. (D) Immunoblot analysis of WCL from P3HR-1 cells cultured in replete or arginine-free media for 5 days, with or without 100 mg/ml acyclovir (ACV) to block EBV lytic replication. (E) Histidine restriction effects on EBV lytic reactivation. Immunoblot analysis of WCL from P3HR-1, MUTU I or EB3 Burkitt cells cultured in media for three days with the indicated histidine level, where 100% histidine refers to the typical RPMI histidine concentration (97μM).

**Figure S2**

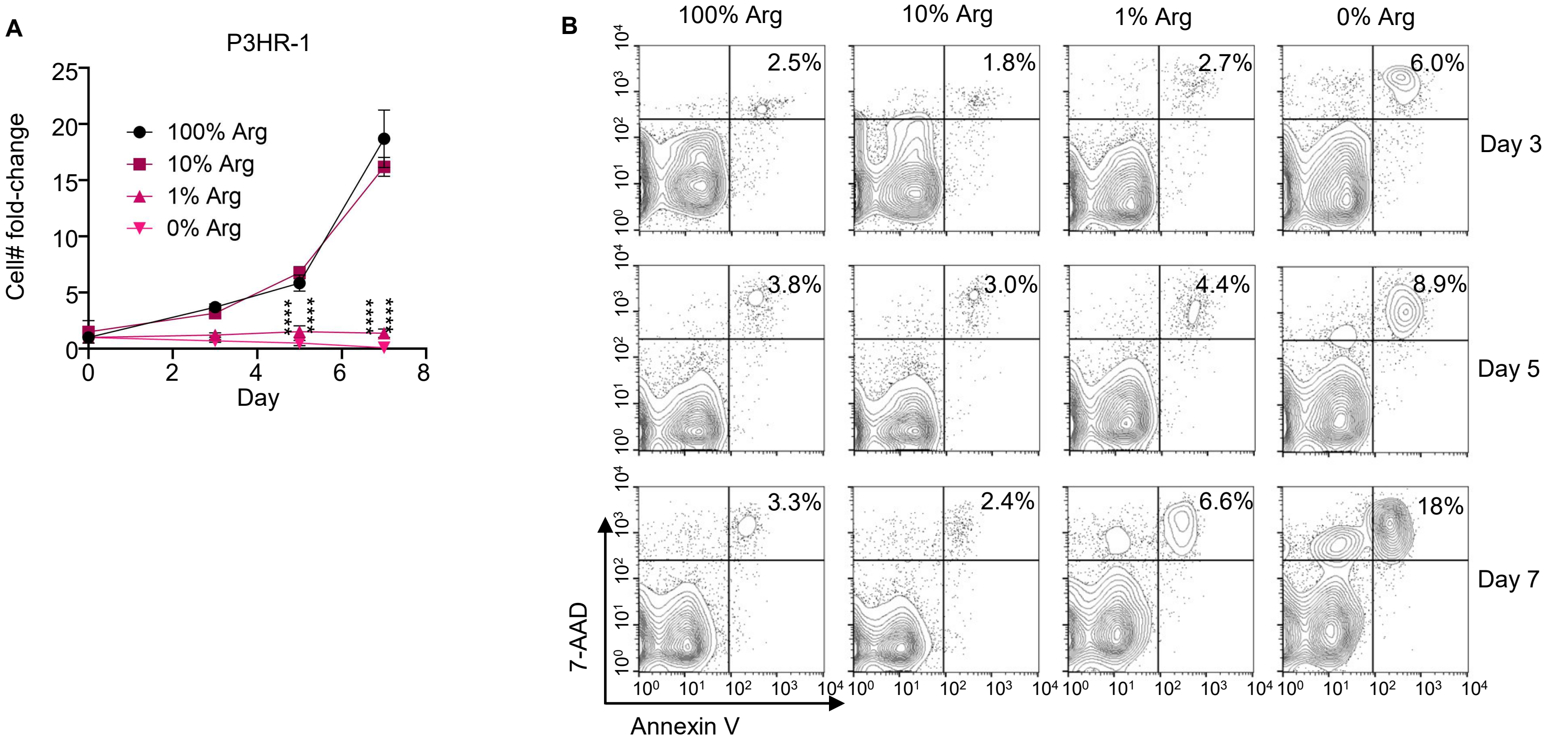
A
P3HR-1
Cell# fold-change
Day
100% Arg
10% Arg
1% Arg
0% Arg
****
****
****
****
B
100% Arg
10% Arg
1% Arg
0% Arg
2.5%
1.8%
2.7%
6.0%
Day 3
3.8%
3.0%
4.4%
8.9%
Day 5
3.3%
2.4%
6.6%
18%
Day 7
7-AAD
Annexin V

**Figure S2. Analysis of arginine restriction effects on P3HR-1 Burkitt growth and survival.**
(A) Analysis of arginine restriction effects on P3HR-1 Burkitt B cell proliferation. Growth curves of P3HR-1 cells cultured in media with the indicated arginine (Arg) concentration. 100% arginine refers to the RPMI arginine concentration of 115 μM. Mean ± SD values from n = 4 replicates are shown. (B) FACS analysis of cells in (A).

**Figure S3**

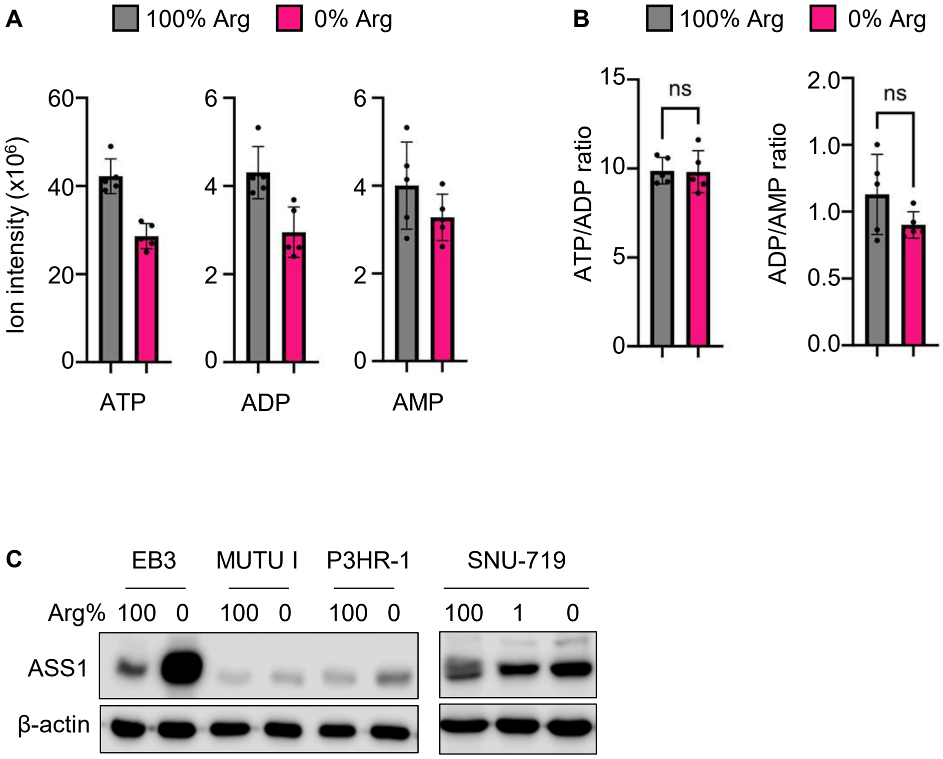
A
100% Arg
0% Arg
Ion intensity (x10^6)
60
40
20
0
6
4
2
0
ATP
ADP
AMP
B
100% Arg
0% Arg
ATP/ADP ratio
15
10
5
0
ns
ADP/AMP ratio
2.0
1.0
1.0
0.5
0.0
ns
C
EB3
MUTU I
P3HR-1
SNU-719
Arg%
100
0
100
0
100
0
100
1
0
ASS1
β-actin

**Figure S3. Arginine restriction effects on ATP, ADP and AMP, related to Figure 3.**
(A) Analysis of arginine restriction effects on adenosine nucleotide levels, from LC/MS metabolomic analysis of EB3 cells grown in replete (100% Arg) vs arginine free (0% Arg) media, as in Figure 3A. (B) Analysis of arginine restriction effects on the ATP/ADP or ADP/AMP ratios, from the LC/MS analysis presented in Figure 3A. (C) Analysis of arginine restriction effects on ASS1 expression. Immunoblot analysis of WCL from the indicated Burkitt cells cultured in arginine replete vs free RPMI for 5 days.

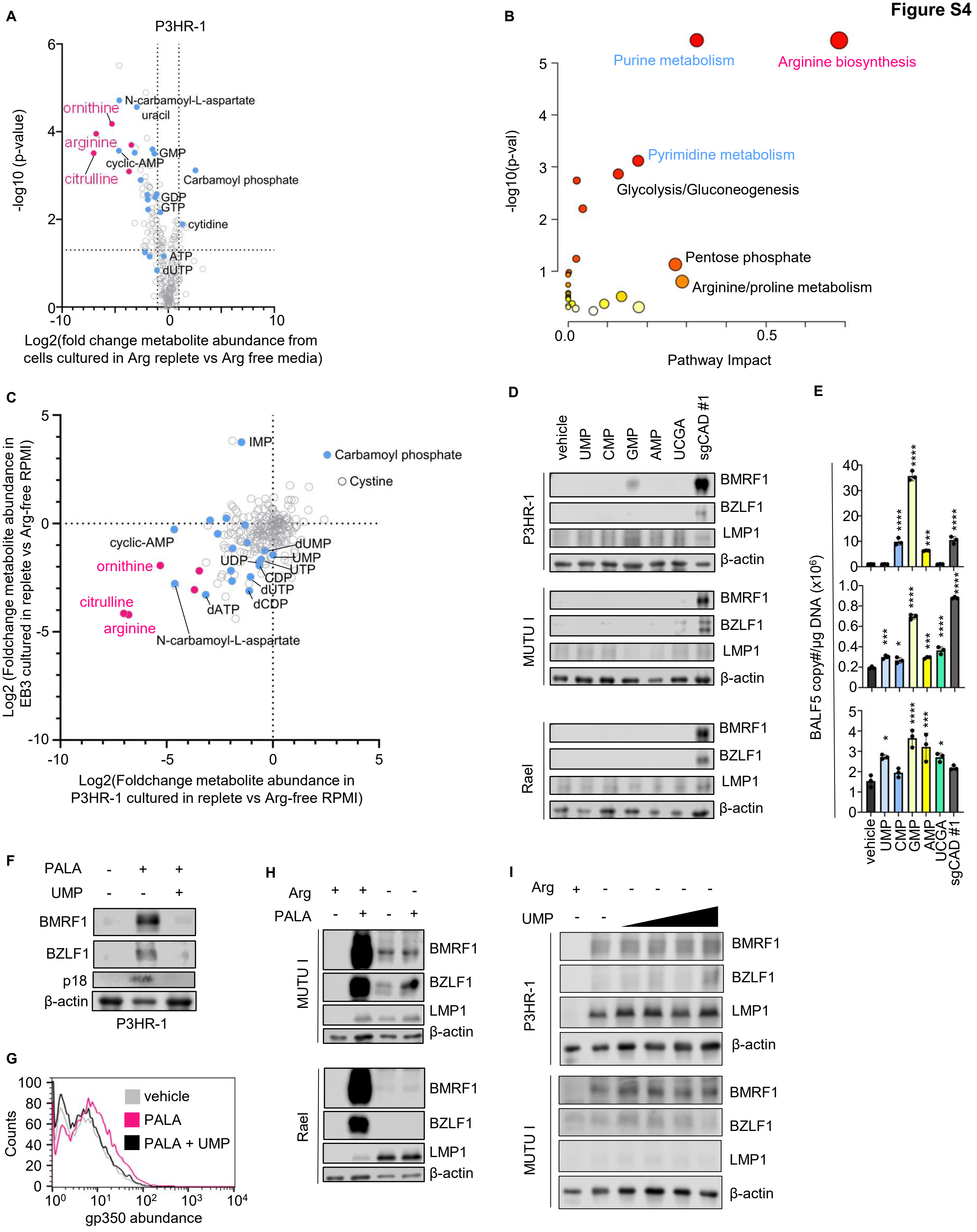
A
P3HR-1
-log10 (p-value)
Log2(fold change metabolite abundance from cells cultured in Arg replete vs Arg free media)
ornithine
arginine
citrulline
N-carbamoyl-L-aspartate
uracil
GMP
cyclic-AMP
Carbamoyl phosphate
GDP
GTP
cytidine
ATP
dUTP
B
-log10(p-val)
Pathway Impact
Purine metabolism
Arginine biosynthesis
Pyrimidine metabolism
Glycolysis/Gluconeogenesis
Pentose phosphate
Arginine/proline metabolism
C
Log2 (Foldchange metabolite abundance in EB3 cultured in replete vs Arg-free RPMI)
Log2(Foldchange metabolite abundance in P3HR-1 cultured in replete vs Arg-free RPMI)
IMP
Carbamoyl phosphate
Cystine
cyclic-AMP
ornithine
citrulline
arginine
N-carbamoyl-L-aspartate
dUMP
UMP
UDP
UTP
CDP
dUTP
dCDP
dATP
D
vehicle
UMP
CMP
GMP
AMP
UCGA
sgCAD #1
P3HR-1
MUTU I
Rael
BMRF1
BZLF1
LMP1
β-actin
E
BALF5 copy#/µg DNA (x10^6)
F
PALA
UMP
BMRF1
BZLF1
p18
β-actin
P3HR-1
G
Counts
gp350 abundance
vehicle
PALA
PALA + UMP
H
Arg
PALA
MUTU I
Rael
BMRF1
BZLF1
LMP1
β-actin
I
Arg
UMP
P3HR-1
MUTU I
BMRF1
BZLF1
LMP1
β-actin

**Figure S4.** ***De novo*** **pyrimidine biosynthesis maintains EBV latency, related to Figure 4.**
(A) Metabolic analysis of P3HR-1 cells cultured in arginine-free vs replete RPMI for 5 days. Shown are foldchanges of metabolite abundance and p-values, calculated from n=3 replicates. Arginine related metabolites are highlighted in pink; pyrimidine and purine related metabolites are highlighted in blue. Higher foldchange indicate higher metabolite abundance in cells grown in arginine replete than arginine free media. (B) KEGG metabolic pathway analysis of metabolomic data as in (A). Metabolites were selected using a FDR<0.05 cutoff and pathway impact values were computed by MetaboAnalyst 3.0 topological analysis. (C) Volcano plot visualization of log2 foldchange in metabolite abundance from LC/MS analysis of P3HR-1 cells (x-axis) versus of EB3 cells (y-axis) cultured in arginine replete vs free media for 5 days. Arginine cycle metabolites are highlighted in pink; pyrimidine and purine related metabolites are highlighted in blue. (D) Immunoblot analysis of WCL from P3HR-1 cells that were cultured in RPMI supplemented with the indicated nucleotide monophosphates to a total concentration of 50μg/ml. Shown at right as a + control of lysates from CAD depleted cells, taken at 6 days after CAD-targeting sgRNA expression. (E) qPCR analysis of intracellular EBV genome copy number of cells as in (D). Shown are the mean ± SD values from n=3 replicates. (F) Immunoblot analysis of WCL from P3HR-1 cells cultured in media with 250μM PALA and with vehicle versus 50μg/ml UMP for 4 days, as indicated. (G) FACS analysis of plasma membrane gp350 abundance in cells as in (F). (H) Immunoblot analysis of WCL from MUTU I or Rael cells cultured in arginine replete or free media, in the absence or presence of 250μM PALA for 4 days. (I) Immunoblot analysis of WCL from cells cultured in arginine replete or arginine free media with 0μg/ml, 0.5μg/ml, 5μg/ml, 50μg/ml or 500μg/ml UMP supplementation for 5 days. Blots are representative of n=3 replicates.

Figure S5

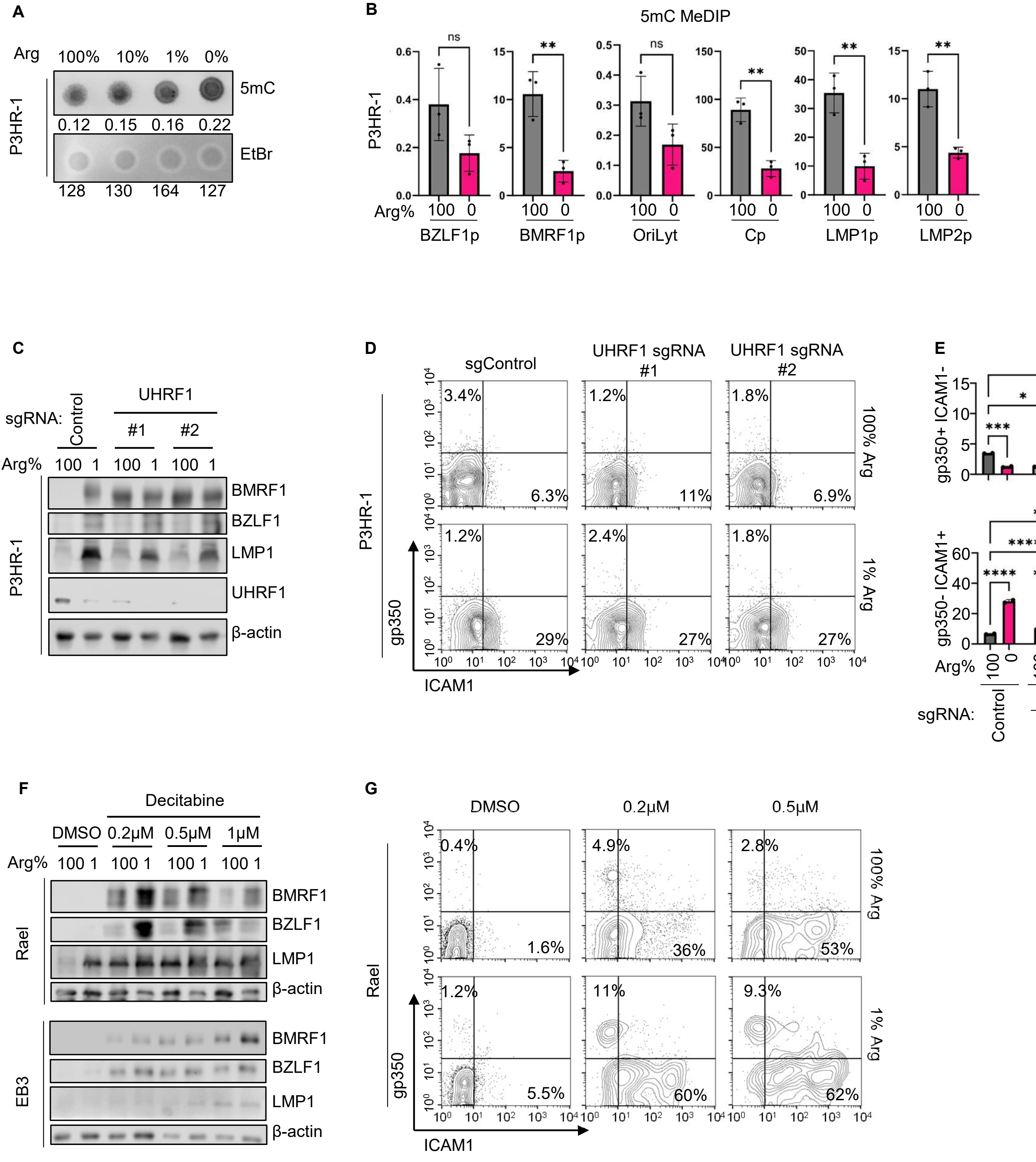

A
Arg 100% 10% 1% 0%
5mC
0.12 0.15 0.16 0.22
EtBr
128 130 164 127
P3HR-1
B
5mC MeDIP
P3HR-1
Arg% 100 0
BZLF1p
BMRF1p
OriLyt
Cp
LMP1p
LMP2p
ns
**
C
UHRF1
sgRNA:
Control
#1
#2
Arg% 100 1 100 1 100 1
BMRF1
BZLF1
LMP1
UHRF1
β-actin
D
sgControl
UHRF1 sgRNA #1
UHRF1 sgRNA #2
100% Arg
1% Arg
gp350
ICAM1
3.4%
6.3%
1.2%
11%
1.8%
6.9%
1.2%
29%
2.4%
27%
1.8%
27%
E
gp350+ ICAM1-
gp350- ICAM1+
Arg%
sgRNA:
Control
UHRF1
F
Decitabine
DMSO 0.2μM 0.5μM 1μM
Arg% 100 1 100 1 100 1 100 1
Rael
EB3
BMRF1
BZLF1
LMP1
β-actin
G
DMSO
0.2μM
0.5μM
Rael
gp350
ICAM1
100% Arg
1% Arg
0.4%
1.6%
4.9%
36%
2.8%
53%
1.2%
5.5%
11%
60%
9.3%
62%

**Figure S5. Arginine restriction and DNA hypomethylation combinatorial effects on EBV reactivation, related to Figure 5.**
(A) 5mC dot blot analysis of DNA extracted from P3HR-1 cells that were cultured in RPMI with the indicated arginine levels for 5 days. Membranes were stained with ethidium bromide as load controls. Each dot contains 500 ng DNA. (B) 5mC MeDIP analysis of chromatin from P3HR-1 cells cultured in arginine free versus replete RPMI media for 5 days. 100 µg/ml acyclovir was added to prevent lytic DNA replication. Mean ± SD values from n = 3 replicates are shown. (C) Immunoblot analysis of WCL from P3HR-1 cells that expressed control or *UHRF1*-targeting sgRNA and were cultured in arginine restricted (1% Arg) vs replete media. Cells were induced to express sgRNA for 3 days and then cultured in the indicated media for 5 days. (D) FACS analysis of plasma membrane gp350 and ICAM-1 expression on cells treated as in (C). (E) Mean ± SEM percentages of gp350+ and ICAM-1+ cells from n=3 replicates from n=2 independent experiments. (F) Immunoblot analysis of WCL from Rael or EB3 cells that were cultured in media containing indicated DCB concentrations and then cultured in 1% Arg or 100% Arg media for 5 days. (G) FACS analysis of plasma membrane gp350 and ICAM-1 levels in Rael cells treated as in (F). Two-way ANOVA or Student's T test was performed, with ****$p < 0.0001$, ***$p < 0.001$, **$p < 0.01$, *$p < 0.05$.
