## supplemental figures and tables for "Arginine Metabolism Supports *De Novo* Pyrimidine Biosynthesis to Block DNA Damage and Maintain Epstein-Barr Virus Latency": Tabel S3 Reagents.docx

| **REAGENT or RESOURCE** | **SOURCE** | **IDENTIFIER** |
| --- | --- | --- |
| **Antibodies** | | |
| Anti-β-actin rat monoclonal antibody | Biolegend | 664802 |
| Anti-LMP1 (S12) mouse monoclonal antibody | Produced from S12 hybridoma cells | N/A |
| Anti-EBNA2 mouse antibody, clone R3 | Millipore Sigma | MABE8 |
| Anti-EBV Ea-D mouse monoclonal antibody (0261): | Santa Cruz Biotechnology | sc-58121 |
| Anti-BZLF1 mouse monoclonal antibody | Santa Cruz Biotechnology | sc-53904 |
| Anti-EBVgP350 (72A1) mouse monoclonal antibody | BioXCell | N/A |
| Epstein Barr Virus p18 Polyclonal Antibody | Invitrogen | PA1-73003 |
| Anti-5-methylcytosine mouse monoclonal  antibody | Abcam | ab10805 |
| Anti-ASS1 rabbit antibody | Millipore Sigma | HPA020896 |
| CAD rabbite polyclonal antibody | Protein Tech | 16617-1-AP |
| DHODH rabbit polyclonal antibody | Protein Tech | 14877-1-AP |
| Anti-UHRF1 Rabbit polyclonal antibody | Diagenode | C15410258 |
| Anti-phospho-ATR (Ser428) Antibody | Cell Signaling Technology | 2853 |
| Anti-phospho-ATM (Ser1981) (D25E5) Rabbit mAb | Cell Signaling Technology | 13050 |
| Anti-phospho-Chk1 (Ser345) (133D3) Rabbit mAb | Cell Signaling Technology | 2348 |
| Anti-phospho-Chk2 (Thr68) (C13C1) Rabbit mAb | Cell Signaling Technology | 2197 |
| Anti-Mouse IgG, HRP-coupled secondary antibody | Cell Signaling Technology | 7076 |
| Anti-Rabbit IgG, HRP-coupled secondary antibody | Cell Signaling Technology | 7074 |
| Anti-Rat IgG, HRP-coupled secondary antibody | Cell Signaling Technology | 7077 |
| IRDye® 800CW Goat anti-Mouse IgG secondary antibody | LI-COR INC | 926-32210 |
| IRDye® 800CW Goat anti-Rat IgG secondary antibody | LI-COR INC | 926-32219 |
| FITC Annexin V antibody | Biolegend | 640945 |
| PE-conjugated Mouse Anti-Human CD54 (ICAM1) | BD Bioscience | 555511 |
| Anti-gp350/220 mouse monoclonal antibody (OT6) | Dr. Jaap Middeldorp | N/A |
| **Cultural media and chemicals** | | |
| DMEM, high glucose, pyruvate | Life Technologies | 11875135 |
| RPMI 1640 Medium | Life Technologies | 11995081 |
| RPMI 1640 Medium for SILAC | Thermo Fisher | A33823 |
| RPMI 1640 medium without amino acids, sodium phosphate | Thermo Fisher | R8999-04A |
| Fetal Bovine Serum, Dialyzed | Thermo Fisher | 26400044 |
| L-Lysine monohydrochloride | Thermo Fisher | A16249.18 |
| L-Arginine | Thermo Fisher | A15738.22 |
| L-Histidine | Millipore Sigma | H8000 |
| L-Phenylalanine | Thermo Fisher | J63925.22 |
| L-Serine | Thermo Fisher | J62187.09 |
| L-Threonine | Millipore Sigma | T8625 |
| L-Tryptophan | Thermo Fisher | J62508.09 |
| L-Tyrosine disodium salt dihydrate | Thermo Fisher | J61770.22 |
| L-Hydroxyproline | ChemScene | CS-W008928 |
| L-Leucine | Millipore Sigma | L8000 |
| L-Isoleucine | Millipore Sigma | I2752 |
| L-Valine | Millipore Sigma | 94619 |
| L-Glutamic acid | Millipore Sigma | G1251 |
| L-Methionine | Millipore Sigma | M5308 |
| L-Asparagine | Millipore Sigma | A4159 |
| L-Aspartic acid | Millipore Sigma | A9256 |
| L-Cystine | Millipore Sigma | C8755 |
| L-Proline | Millipore Sigma | P5607 |
| Glysine | Thermo Fisher | AC120070050 |
| Gibco™ L-Glutamine (200 mM) | Life Technologies | 25030-164 |
| Acyclovir | Millipore Sigma | 114798 |
| Doxycycline hyclate | Millipore Sigma | D9891 |
| Uridine | Millipore Sigma | U3003 |
| Uridine 5′-monophosphate disodium salt | Millipore Sigma | U6375 |
| Guanosine 3′,5′-cyclic monophosphate sodium salt | Millipore Sigma | G6129 |
| Adenosine 5′-monophosphate disodium salt | Millipore Sigma | 01930 |
| Cytidine 5′-monophosphate | Millipore Sigma | C1131 |
| Puromycin Dihydrochloride | Thermo Fisher | A1113803 |
| N-(phosphonacetyl)-L-aspartate (PALA) | NCI | NSC224131 |
| Brequinar (DUP785) | MedChemExpress | HY-108325 |
| Decitabine | Cayman chemical | 11166 |
| **Recombinant DNA** | | |
| pLentiGuide-Puro | Addgene | 52963 |
| pXPR-011 | Addgene | 59702 |
| sgRNA non-targeting ctrl | Broad Institute | BRDN0004438510 |
| pRDA_355 | Addgene | 187159 |
| **Commercial assay kits and reagents** | | |
| Invitrogen PureLink Quick Plasmid Maxiprep Kit | Invitrogen | K210016 |
| DNeasy Blood& Tissue Kit | Qiagen | 69504 |
| Power SYBR Green PCR Master Mix | Applied Biosystems | 4367659 |
| Pierce™ BCA Protein Assay Kit | Thermo Fisher | 23225 |
| MagMeDIP qPCR Kit | diagnode | C02010021 |
| polybrene | Sigma-Aldrich | TR-1003-G |
| UltraPure 10%SDS | Invitrogen | 155553-035 |
| 0.5M EDTA pH8.0 | Invitrogen | 15575-038 |
| Proteinase K | New England Biolabs | P8107S |
| Trypan Blue Solution, 0.4% | Thermo Fisher | 15250061 |
| 7-AAD (7-Aminoactinomycin D) | Thermo Fisher | A1310 |
| ECL chemiluminescence | Millipore Sigma | WBLUF0500 |
| ProLong™ Gold Antifade Mountant with DAPI | Thermo Fisher | P36935 |
| AlbumiNZ™ Low IgG BSA | MPbio | 219989725 |
| BSA, Molecular Biology Grade | NEB | B9000S |
| Cy5® Fast Conjugation Kit | Abcam | ab188288 |
| TransIT-LT1 Transfection Reagent | Mirus | MIR 2306 |
| **Cell lines** | | |
| EBV+ Burkitt lymphoma  P3HR1 ZHT/RHT | Dr. Eric Johannsen | N/A |
| EBV+ Burkitt lymphoma  P3HR1 ZHT/RHT-Cas9 | Ma et al., 2017 | N/A |
| EBV- Burkitt lymphoma  AKATA-Cas9 | Guo et al., 2020 | N/A |
| EBV- Burkitt lymphoma  MUTU I-Cas9 |  |  |
| EBV+ Burkitt lymphoma  MUTU I-Cas9 | Guo et al., 2020 | N/A |
| EBV+ Burkitt lymphoma  Rael-Cas9 | Guo et al., 2020 | N/A |
| EBV+ Burkitt lymphoma  EB3 | ATCC | CCL-85 |
| EBV+ gastric cancer cell  SNU719-Cas9 |  |  |
| 293T | ATCC | CRL-3216 |
| **Software** | | |
| Image Studio Ver 5.5 | Li-COR | https://www.licor.com/bio/image-studio/ |
| GraphPad Prism 7 | GraphPad Software | https://www.graphpad.com/scientific-software/prism/ |
| Flowjo V10 | Flowjo LLC. | <https://www.flowjo.com/> |
| MetaboAnalyst 6.0 | Wishart Research Group | www.Metaboanalyst.ca |
| FIJI ImageJ | Open Source | imagej.net/software/fiji |
| **sgRNAs** | | |
| CAD-sg1 sense | caccgAAGTCAGTAACACACCATCG | |
| CAD-sg1 antisense | aaacCGATGGTGTGTTACTGACTTc | |
| CAD-sg2 sense | caccgCCCAGGAACTCTGTGACAGG | |
| CAD-sg2 antisense | aaacCCTGTCACAGAGTTCCTGGGc | |
| DHODH-sg1 sense | caccgATCTCCCGTGGCCATCAGGT | |
| DHODH-sg1 antisense | aaacACCTGATGGCCACGGGAGATc | |
| DHODH-sg2 sense | caccgCATCTTATAAAGTCCGTCCA | |
| DHODH-sg2 antisense | aaacTGGACGGACTTTATAAGATGc | |
| UHRF1-sg1 sense | caccgACACCCGACTCGCTGACCTG | |
| UHRF1-sg1 antisense | aaacCAGGTCAGCGAGTCGGGTGTc | |
| UHRF1-sg2 sense | caccgTGCTCGGGACACGAACATGG | |
| UHRF1-sg2 antisense | aaacCCATGTTCGTGTCCCGAGCAc | |
| **qPCR primers** | | |
| BALF5_F | GAGCGATCTTGGCAATCTCT | |
| BALF5_R | TGGTCATGGATCTGCTAAACC | |
| LMP1p_F | TCCAGAATTGACGGAAGAGGTT | |
| LMP1p_R | GCCACCGTCTGTCATCGAA | |
| Cp_F | GGCGGGAGAAGGAATAACG | |
| Cp_R | CTTGAGCTCTCTTATTGGCTATAATCC | |
| BZLF1p_F | GCAAGGTGCAATGTTTAGTGAGTT | |
| BZLF1p_R | GCTGGTGCCTTGGCTTTAAAG | |
| BMRF1p_F | ACTGCCCGCTCACCTACAT | |
| BMRF1p_R | CCAGAGCAGAGGCAGGCAGG | |
| OriLyt |  | |

Table S1. Reagents, cell lines, sgRNAs, and qPCR primers used in this study.
